# Input connectivity reveals additional heterogeneity of dopaminergic reinforcement in *Drosophila*

**DOI:** 10.1101/2020.02.19.952648

**Authors:** Nils Otto, Markus W. Pleijzier, Isabel C. Morgan, Amelia J. Edmondson-Stait, Konrad J. Heinz, Ildiko Stark, Georgia Dempsey, Masayoshi Ito, Ishaan Kapoor, Joseph Hsu, Philipp M. Schlegel, Alexander S. Bates, Li Feng, Marta Costa, Kei Ito, Davi D. Bock, Gerald M Rubin, Gregory S. X. E. Jefferis, Scott Waddell

**Affiliations:** Centre for Neural Circuits and Behaviour, The University of Oxford, Tinsley Building, Mansfield Road, Oxford, OX1 3SR, UK; Drosophila Connectomics, Department of Zoology, University of Cambridge, CB2 3EJ, UK; Janelia Research Campus, Howard Hughes Medical Institute, Ashburn, VA 20147, USA; Division of Neurobiology, MRC Laboratory of Molecular Biology, Cambridge CB2 0QH, UK; Institute for Zoology, University of Cologne, Zülpicher Straße 47b, 50674 Cologne, Germany; Department of Neurological Sciences, University of Vermont Medical Center, VT 05405, USA

**Author notes:** Correspondence and Lead Contact Twitter: @_Nils_Otto_, @mwpleijzier, @scottishwaddell, @gsxej, @dddavi.

**Keywords:** Learning and Memory, Extinction, Connectomics, Drosophila, Dopamine, Mushroom Body

## Abstract

Different types of *Drosophila* dopaminergic neurons (DANs) reinforce memories of unique valence and provide state-dependent motivational control [1]. Prior studies suggest that the compartment architecture of the mushroom body (MB) is the relevant resolution for distinct DAN functions [2, 3]. Here we used a recent electron microscope volume of the fly brain [4] to reconstruct the fine anatomy of individual DANs within three MB compartments. We find the 20 DANs of the γ5 compartment, at least some of which provide reward teaching signals, can be clustered into 5 anatomical subtypes that innervate different regions within γ5. Reconstructing 821 upstream neurons reveals input selectivity, supporting the functional relevance of DAN sub-classification. Only one PAM-γ5 DAN subtype γ5(fb) receives direct recurrent input from γ5β’2a mushroom body output neurons (MBONs) and behavioral experiments distinguish a role for these DANs in memory revaluation from those reinforcing sugar memory. Other DAN subtypes receive major, and potentially reinforcing, inputs from putative gustatory interneurons or lateral horn neurons, which can also relay indirect feedback from MBONs. We similarly reconstructed the single aversively reinforcing PPL1-γ1pedc DAN. The γ1pedc DAN inputs mostly differ from those of γ5 DANs and they cluster onto distinct dendritic branches, presumably separating its established roles in aversive reinforcement and appetitive motivation [5, 6]. Tracing also identified neurons that provide broad input to γ5, β’2a and γ1pedc DANs suggesting that distributed DAN populations can be coordinately regulated. These connectomic and behavioral analyses therefore reveal further complexity of dopaminergic reinforcement circuits between and within MB compartments.

**Highlights:**

- Nanoscale anatomy reveals additional subtypes of rewarding dopaminergic neurons.
- Connectomics reveals extensive input specificity to subtypes of dopaminergic neurons.
- Axon morphology implies dopaminergic neurons provide subcompartment-level function.
- Unique dopaminergic subtypes serve aversive memory extinction and sugar learning.

## Results and Discussion

In adult *Drosophila*, anatomically discrete classes of dopaminergic neurons (DANs) innervate adjacent compartments of the mushroom body (MB) lobes [2]. In some cases, it is clear that different combinations of DANs serve discrete roles. However, there are instances where multiple functions have been assigned to DANs that innervate the same compartment. For example, DANs innervating the γ5 compartment reinforce short-term courtship memories, appetitive memories with sugar and they also signal the absence of expected shock, to extinguish aversive memory [7–11]. Similarly, DANs innervating the β’2a compartment have been assigned roles, such as controlling thirst state-dependent water seeking and water memory expression, sugar reinforcement and hunger-dependent modulation of carbon dioxide avoidance [9, 10, 12–14]. Moreover, the individual PPL1-γ1pedc DANs, which innervate the γ1 compartment in both hemispheres are required to reinforce aversive memories with electric shock, high heat and bitter taste, and also provide hunger state-dependent motivational control of sugar memory expression [5, 6, 15, 16]. For an individual DAN to multi-task it must function in different modes.

However, where a compartment is innervated by multiple DANs it is unclear whether different neurons in the population perform discrete functions, and/or whether the group functions together in different modes. Here we used connectomics to investigate the organization of neurons providing input to DANs innervating the γ5, γ1 and β’2a compartments to better understand how valence-specific reinforcement is generated.

### Determining the nanoscale structure of reinforcing dopaminergic neurons

We used a recent EM dataset of a Full Adult Fly Brain (FAFB) [4] to identify, manually trace and reconstruct the nanoscale anatomy of 20 DANs in the Protocerebral Anterior Medial (PAM) cluster whose presynaptic arbors innervate the γ5 compartment (the PAM-γ5 DANs [2]) in the fly’s right brain hemisphere (we also reconstructed 9 PAM-γ5 DANs in the left MB). We then identified 8 right hemisphere PAM-β’2a DAN candidates and reconstructed 6, as well as the 2 Protocerebral Posterior Lateral (PPL)1-γ1pedc DANs that innervate the γ1 compartments of each MB (Figure 1A and S1A-D and Video S1). Metrics of quality control are detailed in STAR Methods.

**Figure 1.**
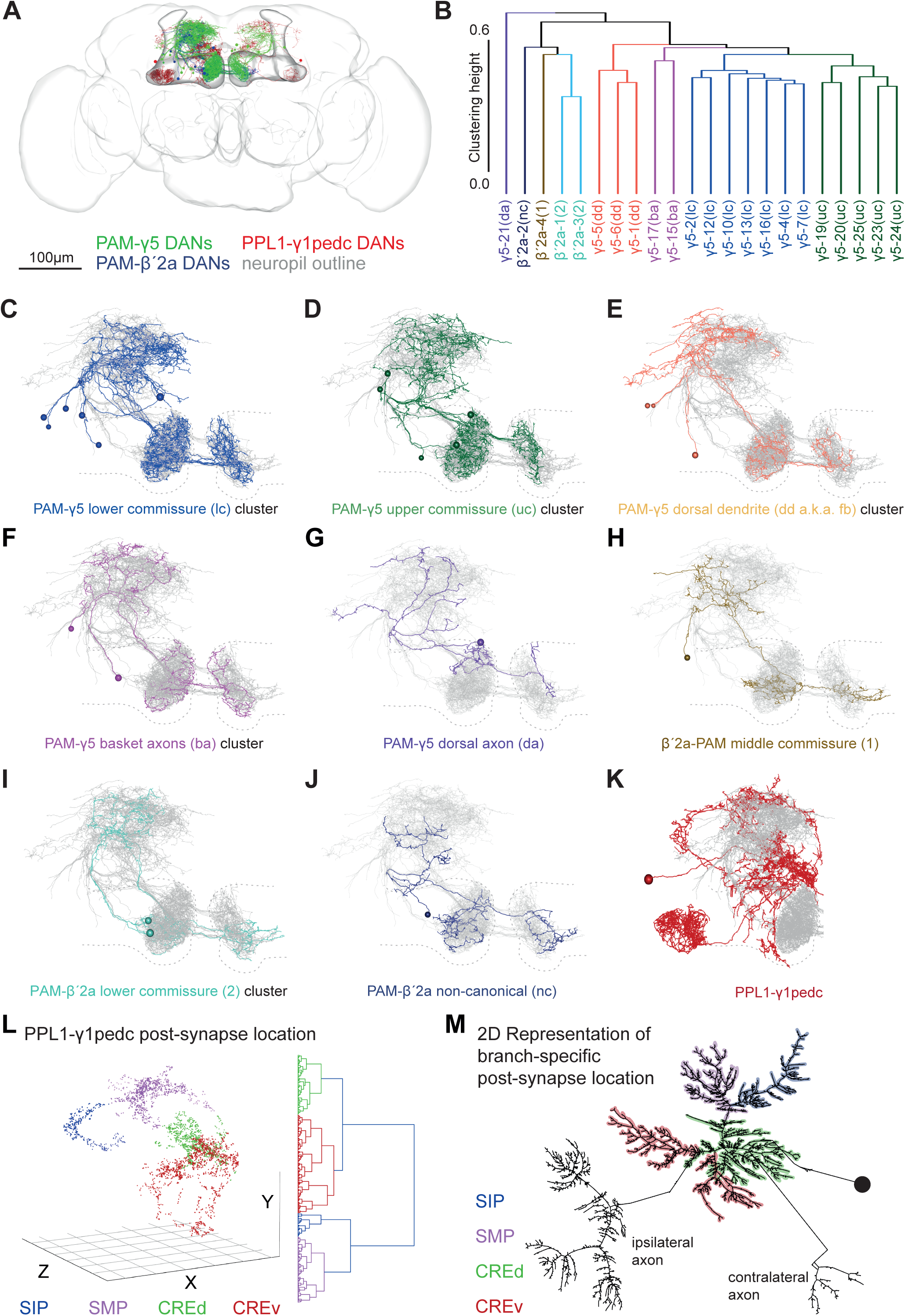
Nanoscale morphology of PAM-γ5, PAM-β’2a and PPL1-γ1pedc DANs reveals new anatomical subtypes and features. (A) Representation of all DANs reconstructed in this study. 20 PAM-γ5 DANs on the fly’s right and 9 on the left, 8 PAM-β’2a DANs on the right, and both left and right PPL1-γ1pedc DANs. The MB and overall brain are outlined. Neuropil reference, Figure S1A. (B) Dendrogram showing hierarchical clustering of PAM DANs by morphology with 5 PAM-γ5 DAN and 3 β’2a DAN clusters. (C-K) Projection views of clustered reconstructed DANs. The morphology of the other traced PAM-γ5, PAM-β’2a DANs are shown in overlap (grey) and the MB neuropil (compare to A) is indicated by a dashed outline. (C) The 7 DANs of the PAM-γ5(lc) cluster (blue). (D) 5 DANs of the PAM-γ5(uc) cluster (green). (E) 3 DANs of the PAM-γ5(dd) cluster (scarlet). Only these neurons receive feedback from MBON-γ5β’2a (Figure 2K) and thus are renamed PAM-γ5(fb). (F) The 2 DANs of the PAM-γ5(ba) cluster (lilac). These DANs occupy the middle commissure. (G) The single PAM-γ5(da) DAN (purple). (H) A PAM-β’2a(1) DAN (ochre) which also occupies the middle commissure. (I) Two PAM-β’2a(2) DANs (turquoise). (J) A non-canonical PAM-β’2(nc) DAN (navy). (K) A PPL1-γ1pedc DAN (maroon). (L) Clustering of PPL1-γ1pedc DAN postsynapses in 3D space generates 4 distinct groups localized in the SIP (superior intermediate protocerebrum), SMP and to both a dorsal and ventral portion of the CRE (CREd and CREv). The optimum number of clusters was determined by the silhouette method, see Figure S1J. (M) Correlation of postsynapse clusters with PPL1-γ1pedc dendritic branches shown on a 2-dimensional dendrogram presented in the graphviz neato layout.

We noticed when reconstructing PAM-γ5 DANs that their dendrites appeared to occupy different areas of the superior medial protocerebrum (SMP) [17], that their somata were connected via 2 main neurite tracts, and that each γ5 DAN had a contralateral projection that crossed the midline of the brain in either an upper, middle, or lower commissure (Figures 1A, C-G and S1A,B and S1E-S1H). We therefore used unbiased anatomical clustering to explore a potential suborganisation of PAM-γ5 DANs (Figure 1B). This analysis grouped PAM-γ5 DANs into 5 discrete clusters comprised of 1-7 neurons (Figures 1C-1G). Importantly, using a different clustering criterion produced an identical result (Figure S1I). Although we did not trace all the fine axonal branches of the PAM-γ5 DANs it was evident that their major presynaptic arbors occupy different areas of the γ5 compartment. We therefore named the PAM-γ5 DAN subtypes according to their defining morphological feature (Figure 1C-G and Video S2).

Four reviewed PAM-β’2a DANs could also be clustered into 3 subgroups with commissure crossed, overall morphology and the region of the compartment that is innervated again being the primary distinguishing features (Figures 1B, 1H-1J and S1C, and S1E-S1H and Video S2). Our tracing also identified a ‘non-canonical’ PAM-β’2a DAN, which mostly innervates β’2a but also extends presynaptic processes into the γ5 compartment (Figure 1J). The dendrites of PAM-β’2a DANs were largely intermingled with those of the PAM-γ5 DANs (Figures 1C-J, Video S1 and S2), consistent with their roles in reinforcing appetitive memories.

Reconstructing the individual right hand PPL1-γ1pedc DAN revealed that its dendrites occupy very different locations to that of the PAM-γ5 and PAM-β’2a DANs (Figure 1K). This suggests that PPL1-γ1pedc receives mostly different input, consistent with it signalling aversive rather than appetitive valence. The three-dimensional structure of the PPL1-γ1pedc DAN dendrite shows four major arbors that extend into discrete locations in the brain (Figures 1L, 1M, S1H and Video S3). Postsynapses also clustered in each of these locations suggesting that PPL1-γ1pedc DANs could receive branch-specific information. This may represent a solution for how a single PPL1-γ1pedc DAN can isolate and prioritize its discrete roles in reinforcement and state-dependent control [5, 6].

### Mapping neuronal inputs onto dopaminergic neurons

We next traced 821 neurons providing input to postsynapses identified in the dendritic arbors of the PAM-γ5, PAM-β’2a and PPL1-γ1pedc DANs (Figure 2A). Since connectivity is dense and manual tracing is very labor-intensive we traced between 45 and 97% of the inputs to the postsynapses annotated on all the reconstructed DANs (Revision Status Table in Materials and Methods). We prioritized upstream tracing so we retrieved a comparable coverage of the inputs to each of the major morphological DAN subtypes. Criteria for sampling and metrics of quality control are again detailed in the Methods.

**Figure 2.**
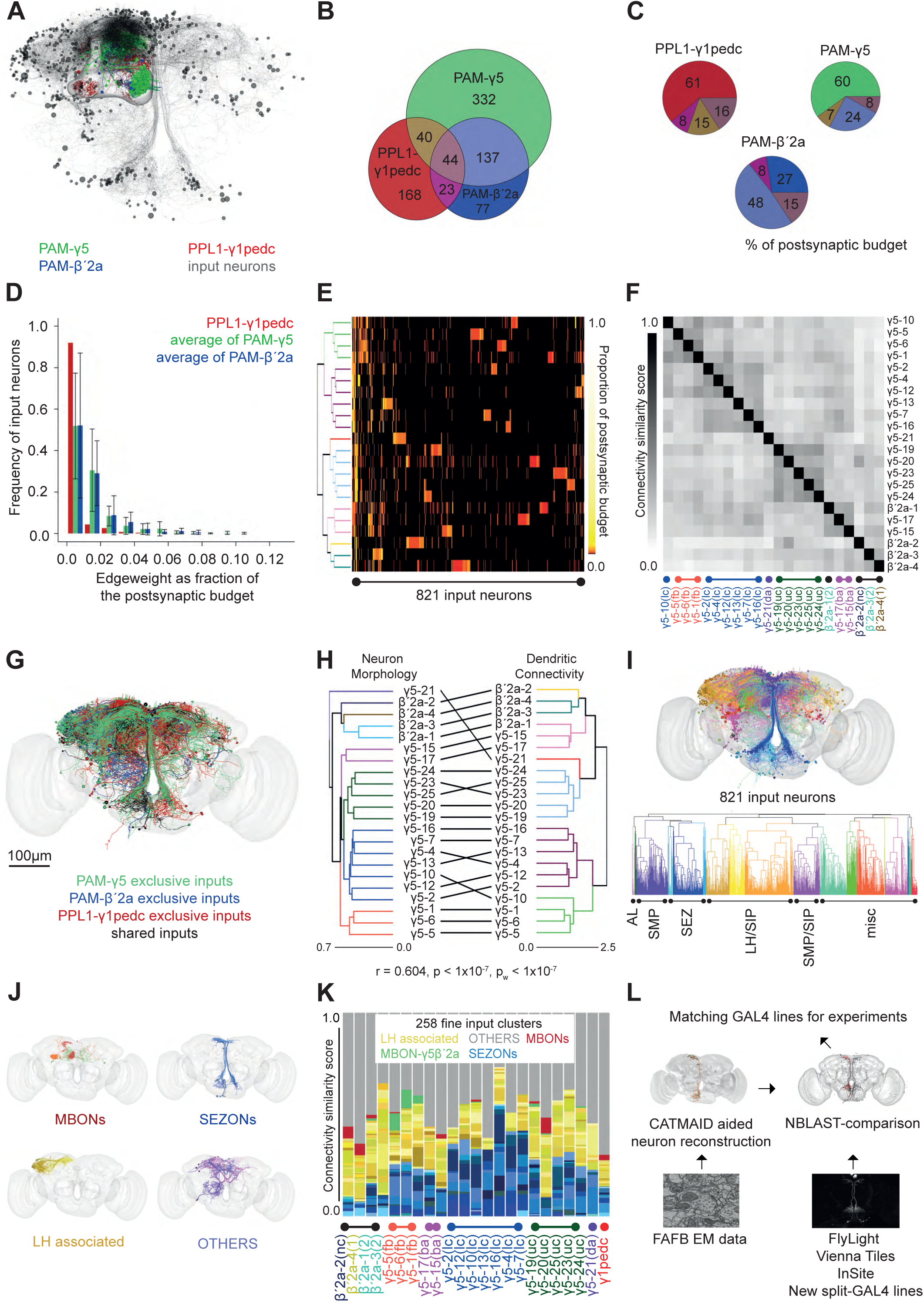
Input specificity to dopaminergic neurons matches anatomical subtypes. (A) Representation of all 821 input neurons to PAM-γ5, PAM-β’2a, and PPL1-γ1pedc DANs identified in this study. Cell bodies (black spheres) and processes (grey). DANs and the MB outline shown for reference (compare to Figure 1A). (B) Venn diagram of unique and common input neurons to the analysed DANs. PPL1-γ1pedc receives largely different input to PAM-γ5 and PAM-β’2a DANs. PAM-γ5 and PAM-β’2a DANs have many common inputs and some are also shared by PPL1-γ1pedc. (C) Pie charts showing percentage of postsynaptic budget occupied by shared and unique input neurons to PAM-γ5, PAM-β’2a, and PPL1-γ1pedc DANs. Percentage of shared inputs across all three groups is 16%, 15% and 8% for PPL1-γ1pedc, PAM-β’2a and PAM-γ5 DANs, respectively. (D) Bar chart showing DANs have many inputs with very low edge weight and each representing a small fraction of their overall postsynaptic budget. Inputs contributing more of the postsynaptic budget (to the right of the graph) are more abundant for PAM-γ5 and PAM-β’2a DANs, PPL1-γ1pedc distribution is strongly left shifted. (E) DANs can be clustered by input connectivity (rows correspond to F). Heatmap shows every DAN has a group of unique input neurons represented by unique blocks in each row. Clustering of DANs mostly depends on lesser number of shared inputs compressed to the left edge of the heatmap. (F) A matrix where DANs are grouped by the similarity of their input connectivity has clear structure, that is significantly more organized than random connectivity (comparison to null-model: p <0.0001, see Methods S1). (G) Representation of traced input neurons labelled using the unique and common input anatomy determined in panel 2B (also see Figure S2A-D). (H) Tanglegram comparing DAN clustering by morphology (from Figure 1B) and clustering by input connectivity (Right of Figure 2E). Connectivity and morphology are not significantly independent of each other (Pearson’s Correlation between the corresponding distance matrices: r=0.604, Mantel Test: p<10^-7^; also pw<10^-7^ within only γ5 or β’2a group. (I) DAN input neurons clustered by morphology. Dendrogram below shows single neurons allocated to 20 major coarse clusters based on soma position and primary neurite tract. Approximate neuropil of origin is indicated: antennal lobe (AL), SMP, SEZ, LH/SIP, SMP/SIP are marked. Many neurons originate from less explored neuropils (misc). (J) Fine clusters of exemplary neurons for the MBON, LHON, SEZON and OTHERS classes of DAN inputs. (K) Barplot showing respective number of MBON, LHON, SEZON and OTHERS inputs to individual DANs, ordered according to cluster identity (Figure 1). In general, PAM-γ5 DANs receive about 35% of their input from SEZONs, and about 20% from LHONs. Only the 3 PAM-γ5(fb) DANs receive significant direct input from MBON-γ5β’2a (green segments). PAM-β’2 DANs receive about 15% from SEZONs and 35% from LHONs. One PAM-β’2a DAN also receives minor direct input from MBON-γ5β’2a. PPL1-γ1pedc DANs receive roughly equal LHON and SEZON input. (L) NBLAST compares CATMAID generated neuronal skeletons from FAFB to neurons labelled in confocal images of GAL4 expression patterns.

Reassuringly, input selectivity of PAM-γ5, PAM-β’2a and PPL1-γ1pedc DANs largely reflected the relative overlap of their dendritic fields. All the DANs receive unique inputs (Figure 2B-C, and S2A-D). However, a greater share of the 553 identified inputs to PAM-γ5 DANs (181) also provided inputs to PAM-β’2a DANs than to the PPL1-γ1pedc DAN (84). Likewise, more of the 281 PAM-β’2a DANs inputs connect to PAM-γ5 DANs (181) than to the PPL1-γ1pedc DAN (67). In contrast more of the 275 traced PPL1-γ1pedc DAN inputs were unique (168) than also contacted PAM-γ5 DANs (84) or PAM-β’2a DANs (67). Lastly, 8% (44) of the traced input neurons synapsed onto all three classes of traced DANs. These common input neurons suggest that the valence specific arms of the DAN system are coordinated under certain contexts. It is however also possible that the different DANs respond in unique ways to the same input neurotransmitters.

Our sampling suggests that despite there being 20 PAM-γ5 DANs and one PPL1-γ1ped DAN, there are only approximately twice as many inputs to all PAM-γ5 DANs compared to the PPL1-γ1pedc DAN (Figure 2B). However, the different DAN types have markedly different weighting of inputs, assuming that synapse number correlates with input strength (Figure 2D and S2E). Whereas the PPL1-γ1pedc DAN receives weakly connected inputs, each input to the PAM-γ5 DANs or PAM-β’2a DANs is more strongly connected and contributes a larger proportion of the individual neuron’s postsynaptic budget. Plotting a connectivity matrix for the most completely traced DANs reveals that certain groups of inputs preferentially synapse onto different PAM-γ5 and PAM-β’2a DANs, demonstrating that all individual PAM-γ5 and PAM-β’2a DANs have an element of input specificity (Figure 2E and S2F).

Nevertheless, a matrix comparing input structure between DANs reveals significant similarities in input between particular groups of PAM-γ5 and PAM-β’2a DANs, which is more organized than random connectivity (Figure 2F and Methods S1). DAN clustering based on dendritic connectivity was identical using two different methods (Figure S2G). Moreover, clustering based on input connectivity correlated well with the prior clustering using full neuron morphology (Figure 2H).

The observed connectivity could be confounded by the incompleteness of reconstruction. However, plotting the synapses identified following standard and extensive review suggests that each iteration of review adds synapses that are evenly distributed across a DAN’s dendritic arbor (Figure S2H). Nevertheless, we tested whether connectivity clustering resulted from unintentional bias in input neuron tracing by repeated clustering following random downsampling of input connectivity from 5-50%. Cluster content remained largely robust in these analyses with DANs clustering within the same groups across 10,000 simulations where synapses were removed (Figure S2I-M). The stability of DAN clustering based on input structure and the similarity of connectivity and morphology clustering suggests that information conveyed by selective input is likely to be maintained in the activity of different DAN subtypes.

We next performed a clustering analysis of the input neurons based on their morphology. In a 3-step approach (Material and Methods) we focused first on soma location and primary neurite layout. This revealed 20 major neuronal clusters (Figure 2I, S2N and Video S4), which could be further decomposed through 2 additional steps into 285 clusters distinguished by the detailed anatomy of smaller neurites. We focused our initial analyses on four classes of input neurons, for which we could postulate a functional role; MBONs, Lateral horn (LH) associated neurons that include Lateral Horn Output Neurons (LHONs), Subesophageal Output Neurons (SEZONs) – potential gustatory projection neurons which ascend from the SEZ, and OTHERS – a variety of single neurons conveying information from other discrete brain areas (Figure 2I-J).

Annotating the DAN clustering with the identity of input neurons revealed that the γ5β’2a MBONs specifically provide feedback (fb) input from the MB to our previously defined PAM-γ5(dd) subtype (green segments in Figure 2K). We therefore renamed the PAM-γ5(dd) neurons as PAM-γ5(fb). Other MBONs (red) also provide selective input to different DANs. In contrast, as a group the SEZONs (blue) and LHONs (yellow) provide input to the PPL1-γ1pedc DAN and all PAM-γ5 and PAM-β’2a DANs, although the relative proportions vary considerably.

Towards assigning functional relevance to the identified input pathways, we first used NBLAST [18] to screen traced neuronal skeletons against a collection of confocal volumes of GAL4 and split-GAL4 lines for those that potentially drive expression in the relevant neurons [19–21] (Figure 2L). This analysis identified >50 driver lines with putative expression in the major SEZONs, LHONs, MBONs and OTHERS groups of DAN inputs.

### Functional analyses of input pathways to DANs

Pairing odor exposure with optogenetic activation of PPL1-γ1pedc or PAM-γ5, PAM-β’2a DANs can produce either aversive or appetitive odor memories, respectively [22]. We therefore assumed that neurons providing significant input to reinforcing neurons should be able to generate similar phenotypes if artificially engaged, instead of the relevant DANs. We therefore combined all the GAL4 driver lines with the red-light activated UAS-CsChrimson [23] optogenetic trigger and screened them for their potential to reinforce olfactory memories by pairing their artificial activation with odor exposure (Figure 3A and S3A). We obtained a broad range of phenotypes. Whereas activation of some GAL4 lines produced robust appetitive odor memories, others produced aversive memories, and some had apparently no consequence. We also correlated the identity of neurons labelled in each GAL4 line with their implanted memory performance and their respective DAN connectivity (Figure 3B, Video S5 and Supplemental Information). These correlations revealed good concordance with the valence of the memories formed by MBONs and SEZONs (see below and Figures 3E-F and S3B for follow-up). We found 3 different LHON types that connect to γ5 DANs and not PPL1-γ1pedc DANs and form appetitive memories; LHON01, LHON02 and LHON-AD1b2 [24, 25] (Figure 3A), and the OTHERS15 line formed aversive memory and preferentially synapses with PPL1-γ1pedc (Figure 3A and B).

**Figure 3.**
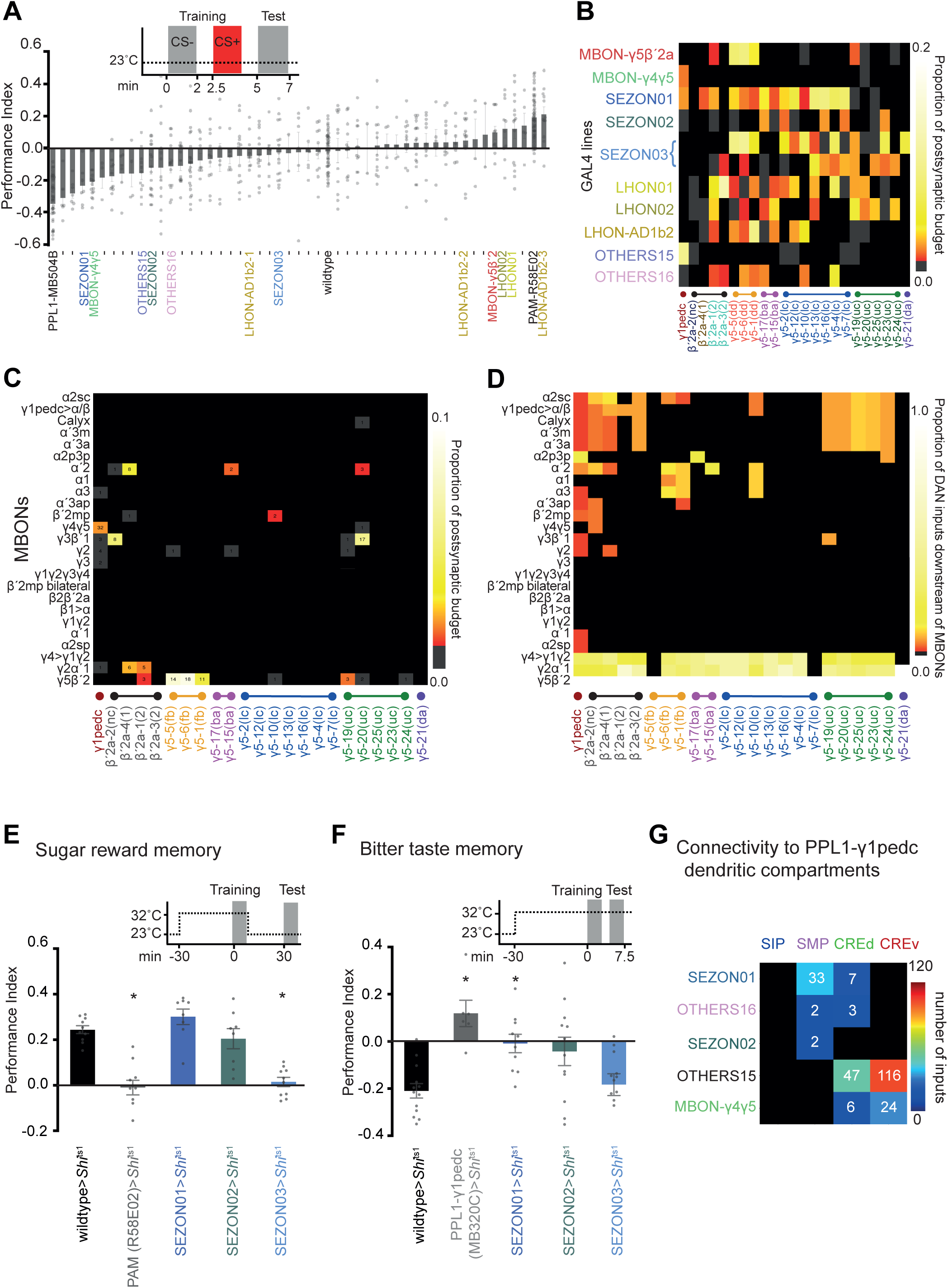
Functional analyses of DAN input neurons. (A) 49 GAL4 driver lines with identified DAN input neurons were used to drive UAS-CsChrimson and screened for memory implantation by pairing neuronal activation with odor exposure. Flies were starved 18-26 h prior to training and tested for immediate memory performance. Lines emphasized in this study (either P.I. >0.1 or <-0.1, or connecting from a neuropil of prior interest are labelled) (Figure S3A, fully labelled version). (B) Connectivity matrix between DANs ordered according to morphological cluster identity and neurons labelled in 10 GAL4 lines, corresponding to 11 input clusters (MBON-γ5β’2a and MBON-γ4γ5, SEZON01-03, LHON01-02, and LHON-AD1b2, and OTHERS15-16). Valence of memory formed is reflected by input connectivity. (C) Direct MBON-DAN connectivity matrix. We identified several MBONs to provide input to specific DANs (note: we traced all inputs to 7 PAM-γ5, and 2 β’2a DANs with extensive review (Figure S2, Methods S1)). Numbers indicate total synapse counts between MBONs and DANs. (D) Adding other traced DAN input neurons creates potential for indirect connectivity between some MBONs and specific subsets of DANs. Indirect connectivity matrix showing the number of DAN input neurons that are downstream of MBONs with at least 3 synapses between each. Columns are normalised by their sum. (E) Olfactory learning with sucrose reinforcement. Schematic, experimental timeline and temperature shift protocol. Blocking neuron output during training abolished 30 min appetitive memory specifically in SEZON03-GAL4; UAS-*Shi*^ts1^ and R58E02-GAL4; UAS-*Shi*^ts1^ flies (p<0.0241 and 0.0089 respectively one-way ANOVA, with Dunnett’s post hoc test, n=10). (F) Olfactory learning with bitter (DEET) reinforcement. Schematic, experimental timeline and temperature shift protocol. Blocking neuron output during training impaired immediate aversive memory in MB320C-, SEZON01- and SEZON02-GAL4; UAS-*Shi*^ts1^, but not in SEZON03; UAS-*Shi*^ts1^ flies. (p<0.0009, 0.0344, and 0.0170 respectively, one-way ANOVA with Dunnett’s post hoc test, n=12). (G) Connectivity matrix to specific branches of the PPL1-γ1pedc dendrite reveals classes of input neurons have branch specificity.

### MBON-DAN connectivity

Two MBONs have previously been described that have a dendrite in the γ5 compartment, MBON-γ5β’2a and MBON-γ4γ5 [3, 11]. In the stimulus replacement screen these MBONs appeared to convey opposite valence. Whereas MBON-γ5β’2a activation formed appetitive memory, MBON-γ4γ5 activation reinforced aversive memory (Figure 3A; and confirmatory 30 min memory experiment in Figure S3B). Connectivity supported these behavioral results. MBON-γ4γ5 is the strongest MBON input to PPL1-γ1pedc, but does not connect to PAM-γ5 or PAM-β’2a DANs, while MBON-γ5β’2a synapses onto PAM-γ5(fb) and a PAM-β’2a DAN but not PPL1-γ1pedc DANs (Figure 3B-C and Figure S3C and S3J).

Our tracing of DAN inputs also identified selective connectivity with a few additional MBONs, some of which were previously unknown (Figure 3C). MBON-*α*’2 was found to synapse onto PAM-β’2a and PAM-γ5(ba) and -γ5(uc) DANs, MBON-β’2mp with PAM-γ5(lc), MBON-γ2*α*’1 with PAM-β’2a, and MBON-γ3β’1 with PAM-β’2a and PAM-γ5(uc). With a few exceptions, the relatively sparse connectivity of MBONs to our traced DANs was largely maintained when we included other traced neurons as potential interneurons between them (Figure 3D). Most notably, the apparent bias of connection of MBON-γ4γ5 to PPL1-γ1pedc and the selectivity of MBON-γ3β’1 remained and new selective clusters to our PAM-γ5(fb), PAM-γ5(uc), and PAM-β’2a groups became apparent. No indirect connections were detected between MBON-γ4γ5 and PPL1-γ1pedc in our dataset. Moreover, only one additional β’2a DAN was seen to be downstream of MBON-γ4γ5 when putative indirect connectivity was included. In contrast, although we found MBON-γ5β’2a to be directly connected to only the PAM-γ5(fb) DANs it is indirectly connected to most of our other traced DANs, and all of the inputs to the unique PAM-γ5(da) neuron come from neurons downstream of MBON-γ5β’2 (Figure 1G). Axo-axonic synapses were frequently observed (>11,000) in the DAN input network. For example, MBONs frequently make reciprocal synapses on the axons of SEZONs and LHONs and the different classes of DAN input neurons are also highly interconnected within cluster (eg. we annotated 317 axo-axonic synapses between SEZON1 neurons).

### SEZON-DAN connectivity

Synthetic activation of the SEZON lines also produced different learning phenotypes. Activation of SEZON01 neurons formed aversive memory and these connect to PPL1-γ1pedc and some PAM-γ5, but not to γ5 DANs reinforcing sugar memory (see below). Stimulating the SEZON03 neurons formed appetitive memory and these SEZONs have synapses onto PAM-γ5 but not PPL1-γ1pedc. SEZON02 neuron activation did not implant significant memory of either valence and appears to connect weakly to all three classes of traced DANs (Figure 3A and B, and S3K).

Despite their specificity, we expect some of our identified GAL4 drivers to express in our traced neurons of interest, and additional similar neurons in a fascicle. For example, a SEZON line could label a collection of ascending neurons representing both tasteful and distasteful gustatory stimuli [26]. Labelling such a mixed population with contradictory value, could explain the inability of SEZON2 to reinforce a memory with clear valence. We therefore used the dominant temperature sensitive UAS-*Shibire*^ts1^ transgene [27] to block neurotransmission from SEZONs during training with sugar or bitter taste reinforcement (Figures 3E-F and S3E-I). We included R58E02-GAL4 as a positive control for sugar memory, which expresses in the majority of PAM DANs [28] and MB320C-GAL4 for bitter learning, because it labels PPL1-γ1pedc [29]. Blocking R58E02 or SEZON03 neurons during training abolished 30 min memory reinforced with sugar but blocking SEZON01 and SEZON02 neurons had no impact. In contrast, blocking MB320C, SEZON01 or SEZON02, but not SEZON03 neurons impaired 30 min memory after bitter learning. These data support a role for SEZONs in relaying positive and negative gustatory valence to DAN subtypes.

Analysis of the location of identified inputs to the PPL1-γ1pedc DAN confirmed that it receives branch-specific information (Figures 3G and S3J-K and Video S6). Whereas all aversively reinforcing input from SEZONs in SEZON01 connect to the arbor in SMP, the OTHERS15 line, which can also produce aversive reinforcement, connect to the PPL1-γ1pedc DAN arbor in the ventral and dorsal crepine (CRE) [17]. MBON input in general and the strong MBON-γ4γ5 inputs are also mostly found on the CREv and CREd branches of PPL1-γ1pedc. Since these branches are the closest to the primary axon, input from other MB compartments may be particularly salient to PPL1-γ1pedc (Figure S3J). Interestingly, the strongest input from MBON-γ5β’2a to the PAM-γ5(fb) DANs are similarly placed on the DAN dendrite (Figure S4A).

### Functional subdivision of PAM-γ5 DANs

Based on prior findings, we hypothesized that PAM-γ5(fb) DANs which receive recurrent feedback from MBON-γ5β’2a would be required for memory revaluation [11] and other PAM-γ5 DANs receiving input from SEZONs labelled by SEZON03 would be required to reinforce sugar memory [9]. Testing this model required locating GAL4 drivers that label subsets of γ5 DANs that at least partially correspond to functionally relevant subtypes. We therefore used the specificity of commissure crossing to select GAL4 drivers, some of which have previously been reported to express in subsets of PAM-γ5 DANs [9]. We reasoned that drivers labelling the lower commissure might express in γ5(fb)-DANs (Figure 1E) and others labelling the upper commissure could include γ5-DANs connected to the appropriate sugar-selective SEZONs. We identified VT006202-GAL4 (Figure 4A), which expresses in γ5-DANs in all 3 commissures, MB315C-(Figure 4B) and 0804-GAL4s (Figure 4C), which express in 8 and 3-5 γ5 DANs respectively in the lower commissure, and 0104-GAL4 (Figure 4D), which only expresses in upper commissure γ5 DANs [2, 9, 21, 30]. We also used genetic intersection with GAL80 to restrict 0104-GAL4 expression to γ5b and β’2m (Figure 4E, [9, 12]).

**Figure 4.**
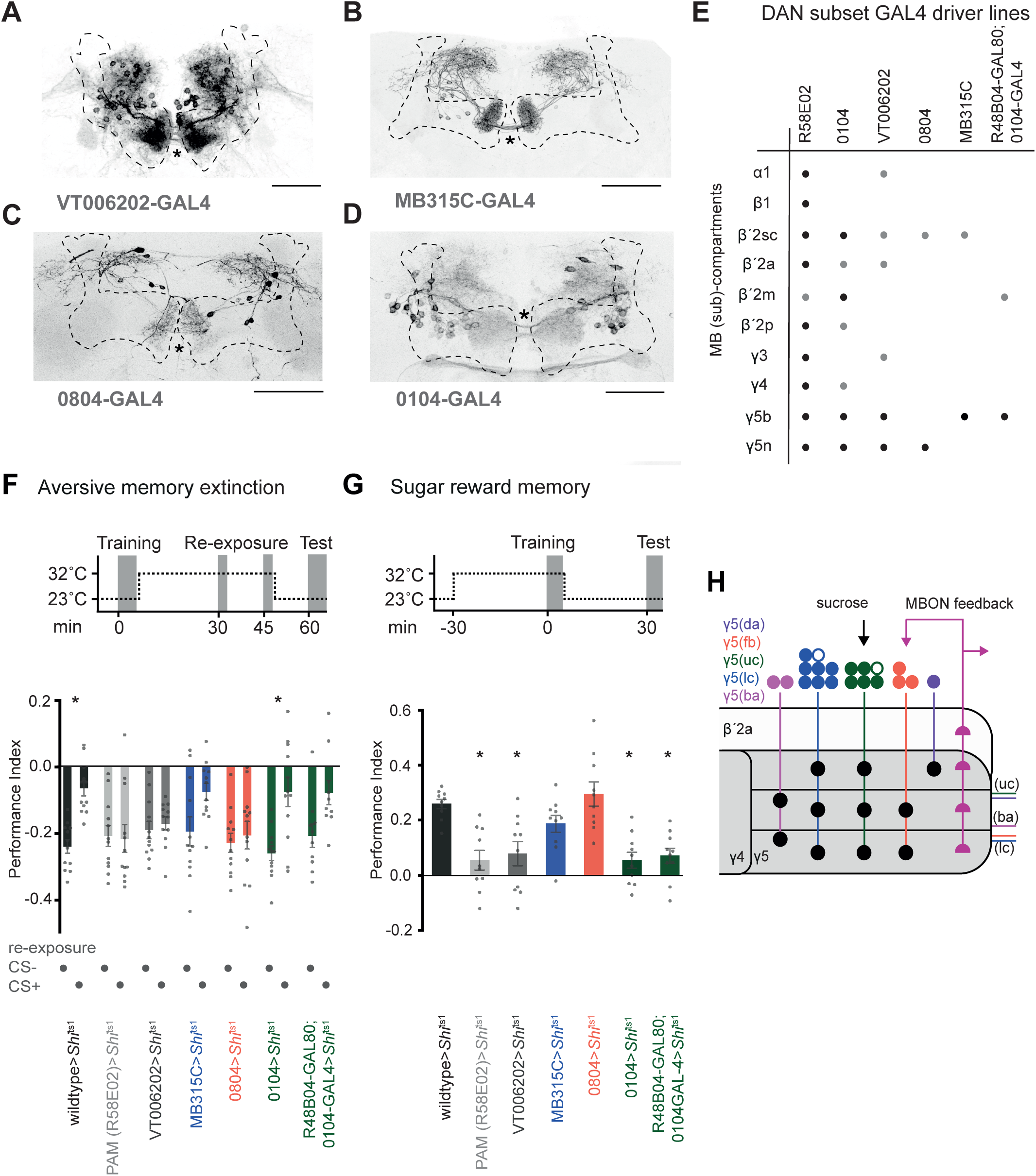
Aversive memory extinction and sugar learning require different subsets of PAM-γ5 DANs. (A) Brain from a VT006202-GAL4; UAS-GFP fly labels all 20 PAM-γ5 DANs and possibly some other PAM DANs (black). 3 commissures are visible (asterisk). Brain co-stained with nc82 antibody (MB is outlined). Scale bar 20µm. (B) MB315C-GAL4; UAS-GFP specifically labels 8 PAM-γ5 DANs per hemisphere that cross the midline in the lower commissure. (C) 0804-GAL4 labels 5 PAM-γ5 DANs per hemisphere, previously ‘γ5 narrow’ that occupy the lower commissure. Scale bar 20µm. (D) 0104-GAL4; UAS-GFP labels PAM-γ5 DANs, previously named ‘γ5 broad’, that cross the midline in the upper and middle commissures. 0104 also labels some other PAM DANs [9]. (E) Table summarizing DAN expression in GAL4 lines used for behavior, modified from [9]. R48B04GAL80 refines the 0104-GAL4 expression [12], shown in the Data S1. (F) Aversive olfactory memory extinction. Schematic, experimental timeline and temperature shift protocol. Blocking neuron output during odor re-exposure impaired memory extinction in R58E02-, VT006202- and 0804-GAL4; UAS-*Shi*^ts1^ but not in MB315C- or 0104-GAL4 +/- GAL80; UAS-*Shi*^ts1^ flies. Asterisks denote p<0.035 (wildtype) and p<0.0176 (0104), one-way ANOVA, with Tukey’s post hoc test, n=10-12). (G). Olfactory learning with sucrose reinforcement. Schematic, experimental timeline and temperature shift protocol. Blocking neuron output during training impaired 30 min appetitive memory in R58E02-, VT006202- and 0104-GAL4 +/- GAL80; UAS-*Shi*^ts1^ but not MB315C- or 0804-GAL4; UAS-*Shi*^ts1^ flies. Asterisks denote p<0.0003 (R58E02), p<0.0004 (0104), p<0.0018 (VT006202), and p<0.0010 (R48B04GAL80; 0104-GAL4); one-way ANOVA, with Dunnett’s post hoc test, n=10). (H) Schematic of input pathways to PAM-γ5 and PAM-β’2a DANs and subtype innervation of the γ5 compartment. Colors correspond to clusters defined in Figure 1. Open circles represent 2 γ5 DANs that were identified but not further analysed.

We next tested whether blocking the neurons labelled in these GAL4s with UAS-*Shi*^ts1^ disrupted aversive memory extinction and/or sugar learning (Figures 4F and G). We again used the broad PAM DAN expressing R58E02-GAL4 as a control. To assay extinction of aversive memory (Figure 4F, Figure S4D), flies were differentially conditioned by pairing one of two odors with electric shock [31]. Then 30 min after training they received two un-reinforced exposures of the previously shock paired odor (CS+) or the other odor (CS-) at 15 min interval [11]. They were then left undisturbed for 15 min before being tested for olfactory preference memory. As previously established, only re-exposure to the CS+ odor diminished memory performance, demonstrating odor-specific memory extinction. We blocked subsets of DANs specifically during odor re-exposure by training flies at permissive 25°C, transferring them to restrictive 32°C immediately after training, then returning them to 25°C after the second odor re-exposure. Blocking R58E02, VT006202 and 0804 neurons abolished memory extinction, whereas memory was still extinguished in flies with blocked MB315C or 0104 neurons (+ and - GAL80). No extinction was observed in any line when flies were re-exposed to the CS-odor after training. These data support a role for the 0804-GAL4 group of lower commissure PAM-γ5(fb) DANs in memory extinction (Figure 4F).

In contrast, when these neurons were selectively blocked only during sugar (sucrose) conditioning [32] (Figure 4G, controls in Figure S4D-E), 30 min memory was impaired in R58E02, VT00602 and 0104 (+ and – GAL80) flies expressing UAS-*Shi*^ts1^, but was unaffected in UAS-*Shi*^ts1^ expressing MB315C and 0804 flies (Figure 4G). These data demonstrate that memory extinction and sucrose reinforcement are dissociable in the γ5 DANs and support the selective role for PAM-γ5(fb) DANs in memory extinction and SEZON connected PAM-γ5 DANs in sucrose reinforcement (Figure 4H). We therefore propose that the other PAM-γ5 DAN subtypes may also serve different reward-related functions.

The morphologically distinct γ5 DANs subtypes innervate different regions of the γ5 compartment where they could depress or potentiate different parts of the KC-MBON network (Figure 1C-G and S4B) [29, 33–35]. We do not currently understand the full relevance of the sub-compartment architecture. However, since the γ-lobe dorsal (γd) KCs carry visual information and the main γ KCs are olfactory [36], connections of these two streams of KCs to γ5 MBONs could be independently modified by PAM-γ5(da) (Figure 1G) and PAM-γ5(dd/fb) DANs (Figure 1E), whose processes are confined to the respective subregions of the γ5 compartment (Figure S4F). DAN stratification may therefore be important for maintaining modality specificity of olfactory memory revaluation [11]. It is interesting to note that larvae only have one DAN per MB compartment [37] and that multiple DANs per compartment is an adult-specific specialization [2]. We expect this expansion reflects the additional behavioral demands of the adult fly and our work here suggests the larger number of DANs in each compartment provides additional functional capacity to the compact anatomy of the adult MB. We propose that distributing inputs onto different branches of its dendrite allows the γ1pedc DAN to multitask [38] but also represents a bottleneck of aversive reinforcement for the fly. Perhaps all bad things are bad or very bad [15, 16]. In contrast, the elaboration and specialization of γ5 DANs may permit the adult fly to individually represent the values of a broad range of rewarding events.

## Acknowledgements

We are grateful to Yaling Huang and Suewei Lin for confocal imaging during COVID-19 shutdown in the UK. We also thank Marion Sillies, Tom Clandinin, Wes Grueber and Wolf Huetteroth for access to confocal stacks. Members of the Waddell and Jefferis groups contributed to discussion throughout this project. This work was largely funded by a Wellcome Collaborative Award (203261/Z/16/Z) to S.W., G.S.X.E.J., D.B. and G.M.R. S.W. is also supported by a Wellcome Principal Research Fellowship (200846/Z/16/Z), the Gatsby Charitable Foundation (GAT3237) and an ERC Advanced Grant (789274).

---

**Figure S1 related to Figure 1.**
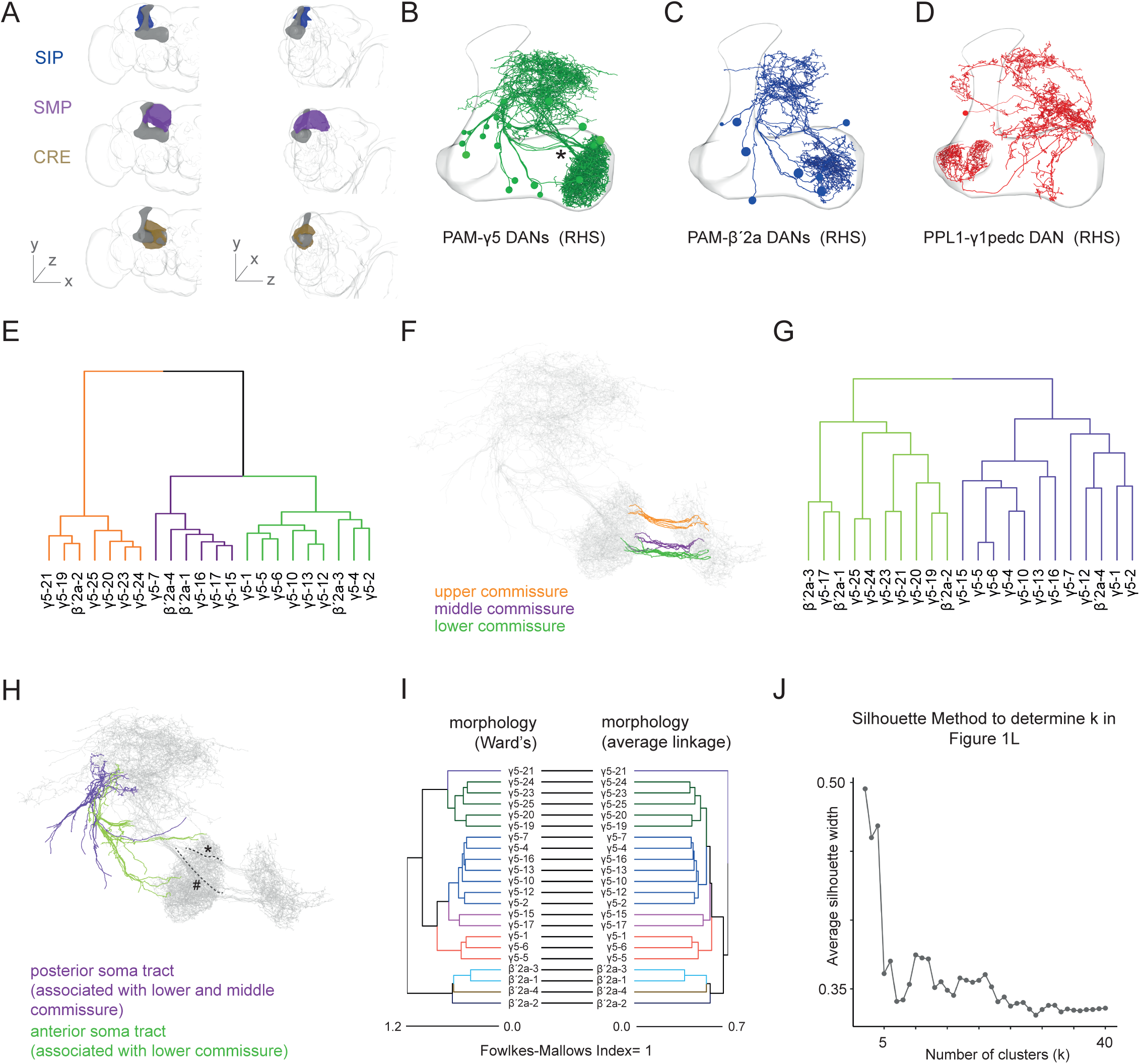
(A) Areas of neuropil providing most input to the DANs in this study as defined in CATMAID and named according to [17]. The superior medial protocerebrum (SMP, purple), superior intermediate protocerebrum (SIP, blue) and crepine (CRE, ochre) are shown from the front (left) and form the midline (right). The MB and whole brain neuropil are depicted in grey. (B-D) Representations of all reviewed DANs in the fly’s right hemisphere. (B) 20 PAM-γ5 DANs. MB outlined, asterisk indicates 2 descending soma tracts. (C) 8 confirmed PAM-β’2a DAN candidates. (D) PPL1-γ1pedc DAN with elaborate dendrite outside the MB. (E) Dendrogram using only the commissure part of the PAM-γ5 and PAM-β’2a DANs reveals 3 morphological clusters. (F) Projection view of the upper (orange), middle (purple) and lower (green) commissure DAN groups from the clustering in (E).Full DAN morphologies are shown in grey. (G) Average linkage clustering using the tracts connecting the somata and dendrites reveals 2 distinct groups. (H) Projection view of the posterior (purple) and anterior (green) soma tracts from clustering in (G). The posterior tract mostly contains lower commissure DANs (dashed line and hash) whereas the anterior tract mostly houses upper commissure DANs (dashed line and asterisk). (I) Tanglegram illustrating comparison of morphological clustering obtained using average linkage criterion (as in Figure 1B) to that retrieved using Ward’s criterion. The clusters are identical, hence Fowlkes-Mallows Index is 1. Note: the between cluster relationship varies slightly. (J) Silhouette method to determine the appropriate number of clusters (k) selected for Ward’s criterion based hierarchical clustering of 3-dimensional positions of PPL1-γ1pedc post synapses. Accuracy drops after 4 clusters and k=4 best reflects the quadripartite structure of the PPL1-γ1pedc dendrite.

**Figure S2 related to Figure 2.**
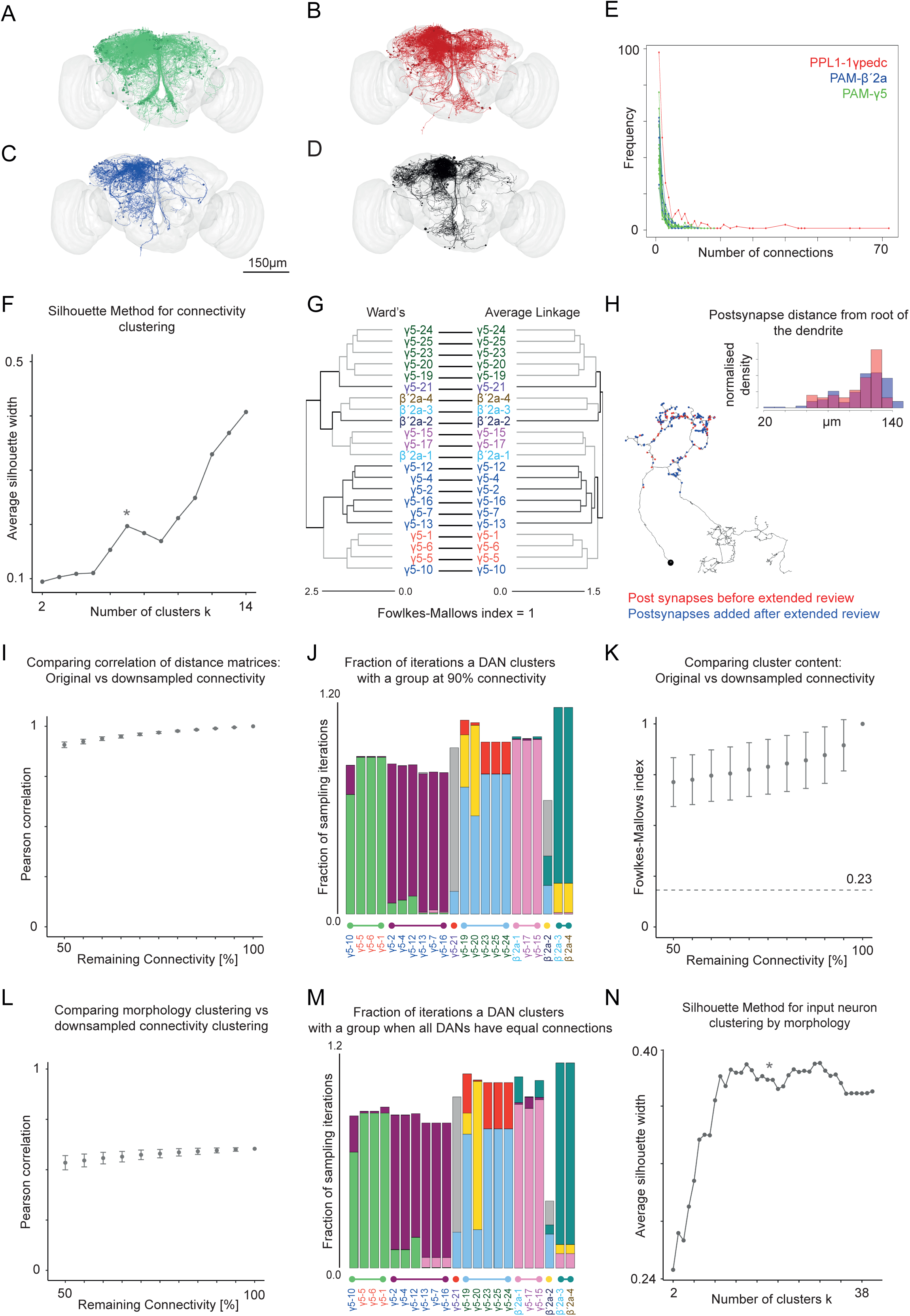
(A-D) Related to Figures 2C and 2G. Representation of input neurons unique to (A) PAM-γ5 DANs (green), (B) PPL1-γ1pedc (red) and (C) PAM-β’2a DANs (blue). (D) Shared inputs (black). Whole brain outlined in grey. (E) Related to Figure 2D. Absolute (non-normalised) edge weight distribution between input neurons and DANs. The frequency at which an input to a given DAN with a certain number of connections occurs is shown as a function of the number of connections. DANs receive many inputs connected with only one or few synapses (skew to left). The long tailed distribution of PPL1-γ1pedc indicates it has some inputs with high absolute edge weights. (F) The Silhouette method was used to determine the appropriate cluster number (k) for connectivity clustering using Ward’s criterion. Due to the high number of unique inputs with low connection number (Figure S2E), a single cluster for each DAN would be optimal. We chose k=7 to reflect the number of morphological DAN subclusters and the local maximum at 7 (asterisk) suggests k=7 is accurate. (G) Tanglegram illustrating comparison of connectivity clustering using Ward’s criterion versus that from average linkage criterion. Identical clusters are formed although the between cluster relationship varies slightly. (H) Comparison of localization of postsynapses created by additional extensive review cycles compared to those identified following normal expert review. Marking the synapses along a γ5 DAN dendrite suggests both classes of synapses to be uniformly distributed. Plotting the distribution of synapses by measuring their positions as distance to the root of the dendritic tree confirms that both the standard and extensively reviewed populations are mixed along the dendrite. These analyses indicate that traced neurons that have undergone standard expert review, but not subjected to extensive review, can be considered to be randomly sampled. (I – L) To test whether input clustering is biased by an incompletely traced input network, we randomly down-sampled the connectivity dataset from 95% – 50% connectivity and compared the clustering obtained from these simulated datasets to the observed connectivity dataset (defined as 100% connectivity). (I) Plot illustrating correlation of connectivity distance matrices for differently down-sampled datasets versus our starting 100% connectivity dataset. The Pearson’s correlation remains >0.9 even at 50% downsampling. The Null model based on randomised cluster allocation has a correlation of 4×10^-4^. (J) To determine whether the DANs cluster in the same groups in the reduced (90%) connectivity dataset we compared their distributions in 10,000 iterations to those in the observed clusters at 100% connectivity. The fraction of times each neuron clusters with a particular group is shown, color coded according to the observed connectivity groups in Figure 2H (Also see cluster similarity analyses in Materials and Methods). Most DANs are reproducibly cluster with the same groups at 90% connectivity and only the γ5-19, γ5-20 and β’2a-2 show any propensity (<40%) to cluster with a different group. (K) The differences in clustering for 10,000 down-sampled datasets at each connectivity density can be individually compared to the observed 100% connectivity dataset using the Fowlkes-Mallows index (FMI). Even 50% connectivity datasets have an FMI (average = 0.71) well above the comparison for a randomised cluster allocation – the null model, FMI = 0.21. (L) Pearson’s Correlation between connectivity and morphology for 100% and down-sampled connectivity datasets shows only subtle changes even at 50% connectivity, indicating that changes in (k) do not greatly alter the correlation between morphological and connectivity groups. (Randomised cluster allocation null model: - 6×10^-4^). (M) To test whether input connectivity clustering results were biased by the varying completeness of DAN post synapse annotation cluster similarity analyses were performed on 10,000 datasets in which all extensively reviewed PAM DANs had their connectivity (on average 216 inputs) down-sampled to the average connectivity of normally reviewed PAM DANs (on average 123 inputs). Only The γ5-20 and β’2a-2 DANs vary in their grouping with β’2a-2 likely to cluster with the γ5(uc) DANs and γ5-20 clustering alone. Clustering of all other DANs is stable indicating that the conclusions of this study do not result from biased completeness of tracing. (N) The Silhouette method was used to determine the accuracy of (k) for clustering based on input structure. Beyond 10 clusters the average silhouette width is high so taken with evaluation of cell body locations k = 20 was deemed to be accurate.

**Figure S3 related to Figure 3.**
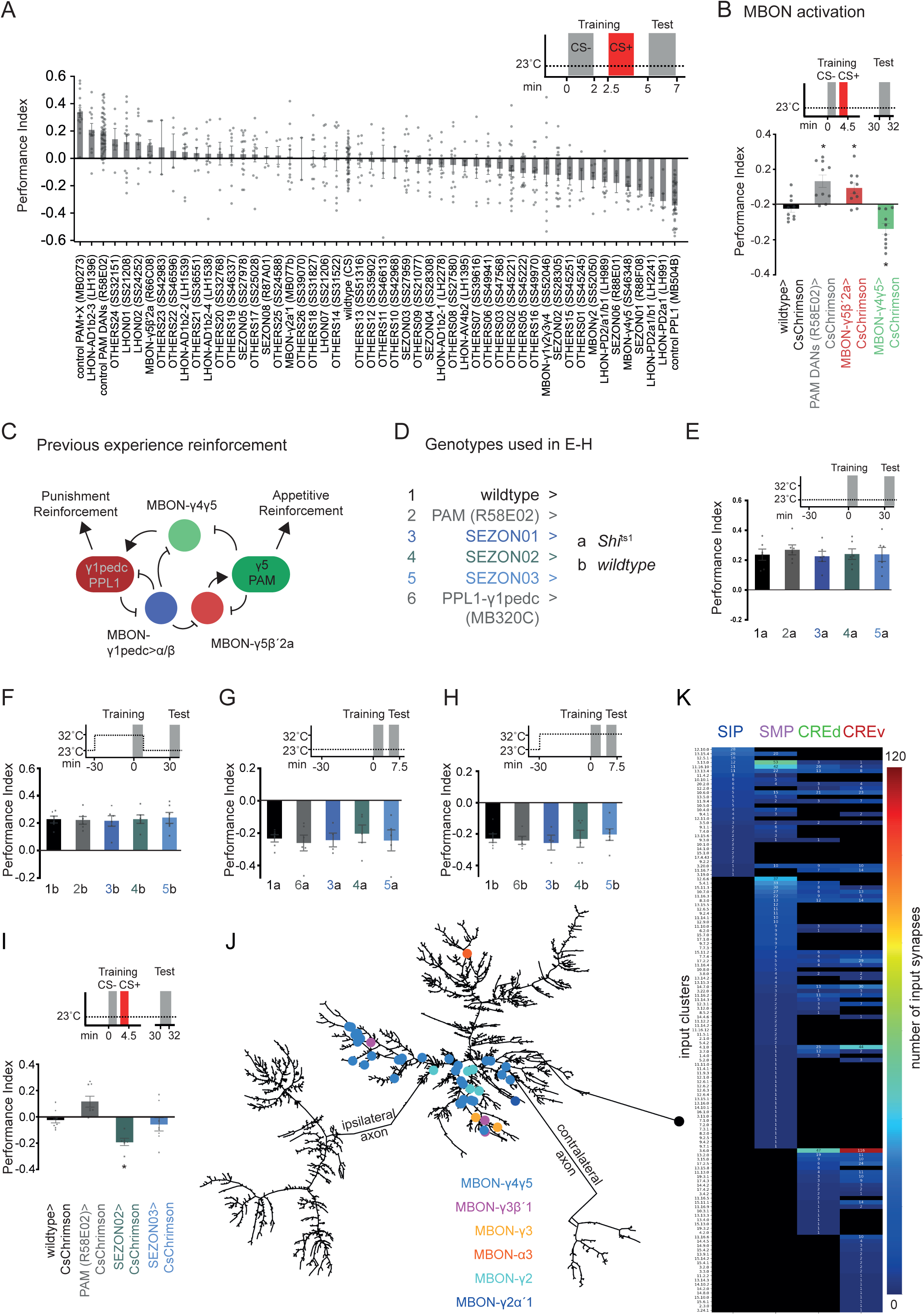
A) Detail of screening results in Figure 3A. (B) MBON activation paired with odor exposure can write memories. Following identification in the screen, MBON-γ5β’2a and MBON-γ4γ5 were retested with 30 min memory being assessed. Activation of MBON-γ5β’2a formed weak positive memory (p<0.0247), whereas MBON-γ4γ5 activation formed aversive memory (p<0.026). PAM DANs (p<0.0021) (PAM DANs). Asterisks denote significance, one-way ANOVA, with Dunnett’s post hoc test for multiple comparisons, n=10. (C) Previous experience reinforcement network for MBON-γ5β’2a and MBON-γ4γ5. Both MBONs receive dendritic input in the γ5 compartment but reinforce memories of opposite valence. MBON-γ4γ5 provides cross-compartmental feedback innervating PPL1-γ1pedc but not PAM-γ5 DANs. MBON-γ5β’2a provides feedback to the same compartment γ5 DANs and does not connect to PPL1-γ1pedc. MBON-γ1pedc>*α*β (MVP2) is also part of the network and provides feed-forward inhibition [38]. (D) Key for groups shown in panels E-I. (E-I) Related to Figure 3 E and F. (E-H) Temperature and genetic controls for *Shi*^ts^ block during training with sucrose (E and F) and DEET reinforcement (G and H). (I) SEZON activation paired with odor followed by testing 30 min memory replicates the screening phenotypes. SEZON02 produces aversive memories (p<0.0245, one-way ANOVA, with Dunnett’s post hoc test for multiple comparisons, n=10). (J) Dendrogram of the dendrite of the PPL1-γ1pedc DAN showing upstream MBON presynapses (neato layout - graphviz). All MBON input goes to the CRE branches closest to the axon. MBON-γ4γ5 provides the strongest MBON input. (K) Relates to Figure 3G. Clustered input neurons provide selective input to specific branches of the PPL1-γ1pedc DAN dendritic tree.

**Figure S4 related to Figure 4.**
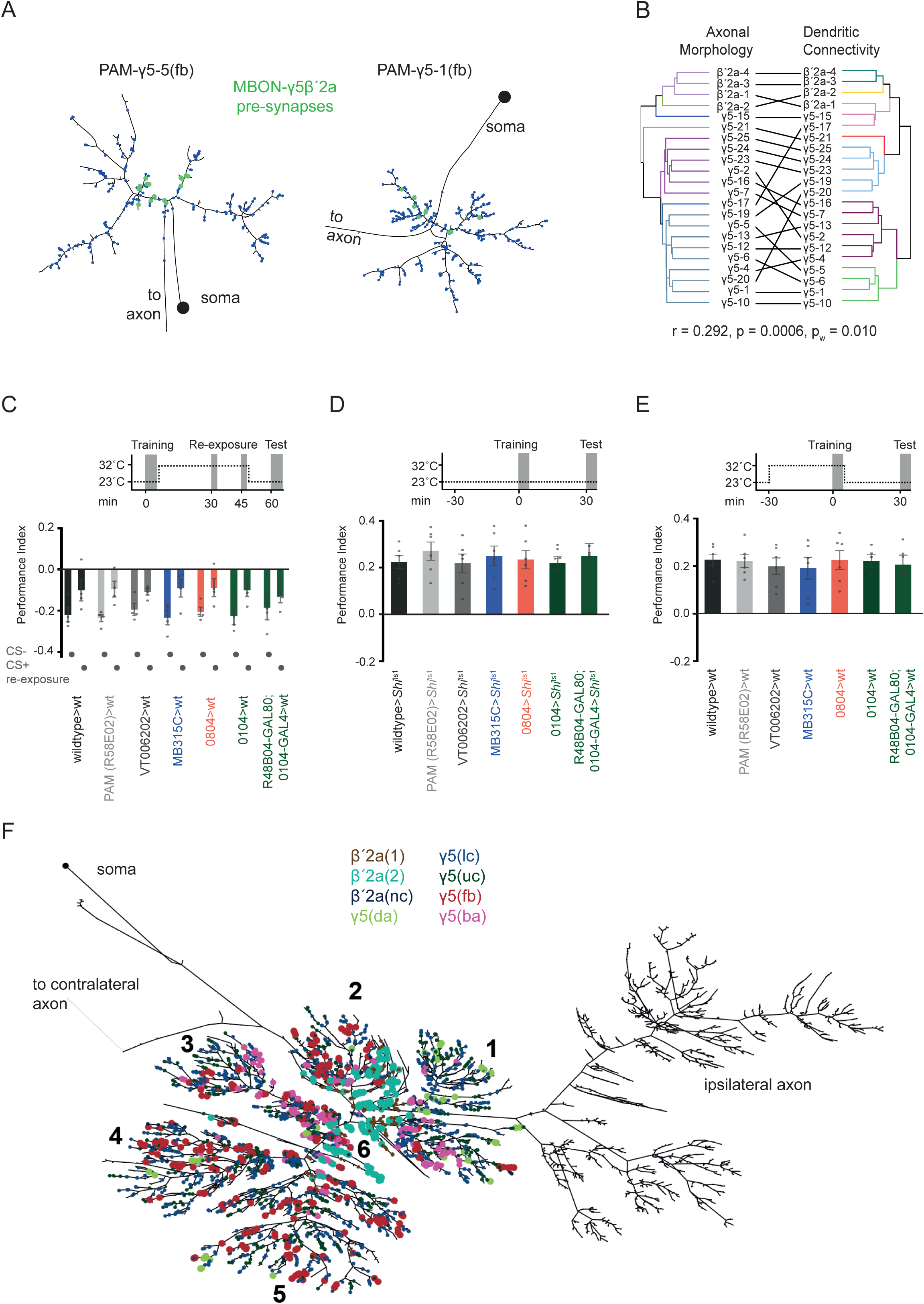
(A) Dendrograms of two PAM-γ5(fb) DANs with MBON-γ5β’2a synaptic inputs shown in green (neato layout - graphviz). MBON-γ5β’2a inputs localize close to the root of the dendrite (left) and on a single branch (right). MBON-γ5β’2a provides the largest fraction of the postsynaptic budget of γ5(fb) DANs contributed by a single neuron. (B) Tanglegram showing the correlation between morphological clustering of PAM DAN axons and dendritic connectivity clustering (Pearson’s Correlation: r=0.292; p=0.0006, pw=0.01 Mantel test). (C) Corresponds to Figure 4F. Heterozygous genetic driver controls show extinction learning (n=4). (D) Corresponds to Figure 4G. Permissive temperature control; no defects in 30 min sugar memory were observed (n=6). (E) Corresponds to Figure 4G. Genetic driver controls for 30 min sugar memory (n=5-6). (F) Dendrogram of MBON-γ5β’2a (neato layout - graphviz) with marked MBON postsynapses. Colors correspond to DAN morphology clusters providing the closest presynapse to the MBON postsynapses. We allowed a 2 µm radius around the respective postsynapse to account for dopamine diffusion and assign DAN presynapses. Dendritic branches are numbered 1-6 where 1, 2, and 6 are closest to the root of the dendrite. PAM-γ5(fb), -γ5(ba), -γ5(da), and -β’2a (2) DANs may influence specific branches, with the γ5(da) and γ5(fb) DANs connect exclusive branches and the γ5ba and β’2a (2) inputs localizing very close to the root of the dendrite. Morphologically distinct PAM DANs that receive different inputs might therefore modulate specific branches of the MBON-γ5β’2a dendrite.

**Video S1 related to Figure 1A**

3D representations of all canonical RHS DAN skeletons traced to identification in this project within the MB neuropil. PPL1-γ1pedc DAN (red, n=1), PAM-γ5 (green, n=20), PAM-β’2a (blue, n=8). Note: for visibility the LHS axonal projections are omitted and only the proximal commissural axons are retained.

**Video S2 related to Figure 1C-1K**

3D representations of skeletons of newly identified subgroups of PAM DANs. Axons from the different subgroups innervate distinct areas in the γ5 and β’2a compartments and their dendrites occupy unique areas in the SMP. Note: for visibility the LHS axonal projections are omitted and only the proximal commissural axons are retained.

**Video S3 related to Figure 1M**

3D representation of the traced RHS PPL1-γ1pedc DAN skeleton within the MB neuropil. The dendritic arbors are labelled by the different areas of neuropil in which they receive synaptic input: SIP (blue), SMP(violet), dorsal CRE (green), ventral CRE (red). The ipsilateral axon, the soma and soma tract are grey. Video V6 details locations of input from MBONs.

**Video S4 related to Figure 2**

3D representations of the skeletons of 821 neurons that provide input to PPL1-γ1pedc, PAM-γ5 and PAM-β’2a DANs. Neurons are shown within the whole brain neuropil color coded according to the first step of coarse clustering (see Star methods for detail). Note: SEZONs are shades of blue, LH-related neurons are shades of yellow, and OTHERS confined to the superior part of the brain are in shades of violet.

**Video S5 related to Figure 3**

3D representation of the skeletons of the fine clusters of input neurons that are studied in detail in this study. Colors correspond to those in Figure 3: LHON01, LHON02, and LHON-AD1b2 neurons are shown in shades of yellow, MBON-γ4γ5s in green, the RHS MBON-γ5β’2a is coral, SEZONs are shades of blue, and OTHERS are shades of violet.

**Video S6 related to Figure 1M and Figure S3K**

3D representation of the skeleton of the RHS PPL1-γ1pedc DAN within the MB neuropil. The dendritic arbors are labelled by the different areas of neuropil in which they receive synaptic input: SIP (blue), SMP (purple), dorsal CRE (green), ventral CRE (red). The ipsilateral axon, soma and soma tract are grey. The dendritic location of input synapses from MBONs are indicated, with most targeting the CRE portion of the dendritic field. MBON-γ4γ5 (light blue, 24xCREv + 6xCREd), MBON-γ2 (turquoise, 5xCREd), MBON γ3β’1 (magenta, 2xCREd + 1xCREv), MBON-γ3 (orange, 2xCREv), MBON-γ2α’1 (dark blue, 1xCREd), MBON-a3 (red, 1x SMP).

## Star Methods

### RESOURCE AVAILABILITY

#### Lead Contact

Further information and requests for resources and reagents should be directed to and will be fulfilled by the Lead Contact, Scott Waddell.

#### Materials Availability

The new split-GAL4 *Drosophila* lines described in this study were produced by Masayoshi Ito. They are available on request from the Lead Contact and will be sent from the Waddell lab or from the Janelia Research Campus, via K. Ito and G. M. Rubin.

#### Data and Code Availability

The datasets and code used for analyses in R and Python are mostly available through public repositories as indicated in this Methods section of the manuscript. Any other code is available on request and without restriction. Neuronal morphologies and connectivity data will be publicly available through www.virtualflybrain.org and neuronal skeletons can be requested from the authors in .swc format.

### KEY RESOURCES TABLE

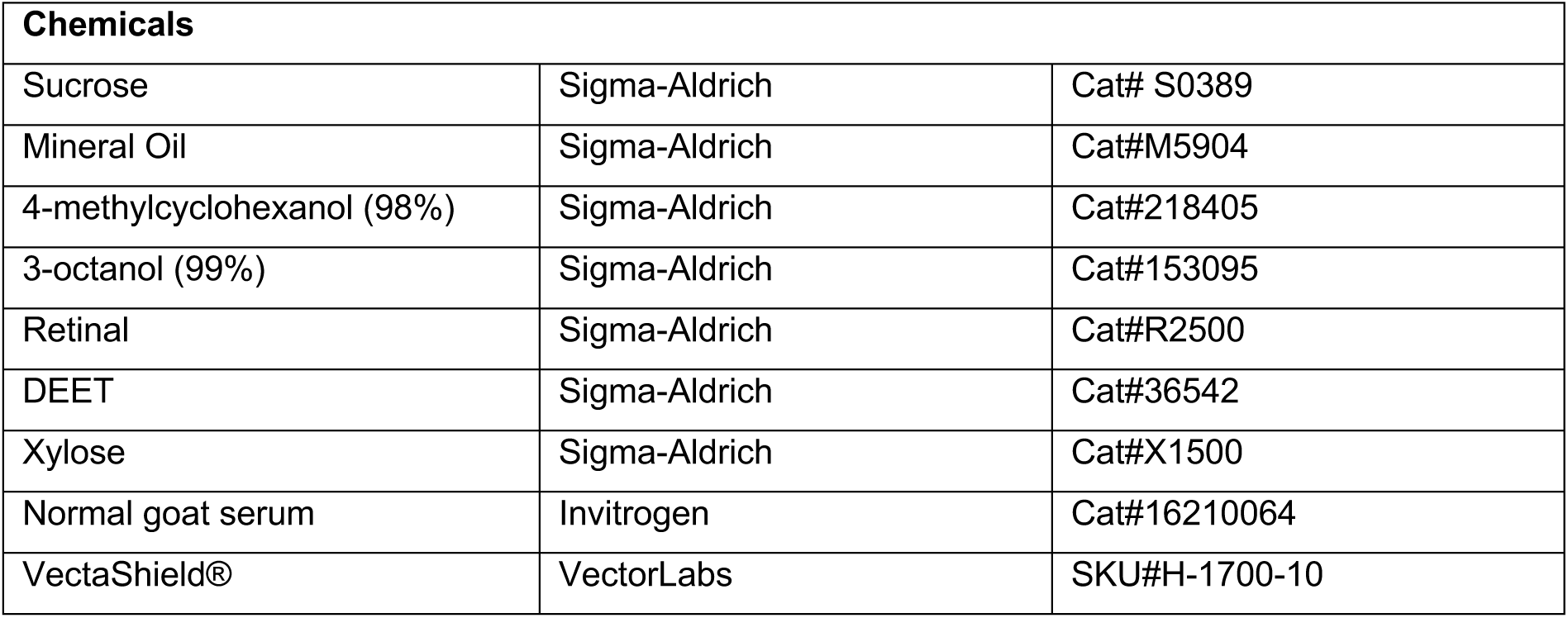
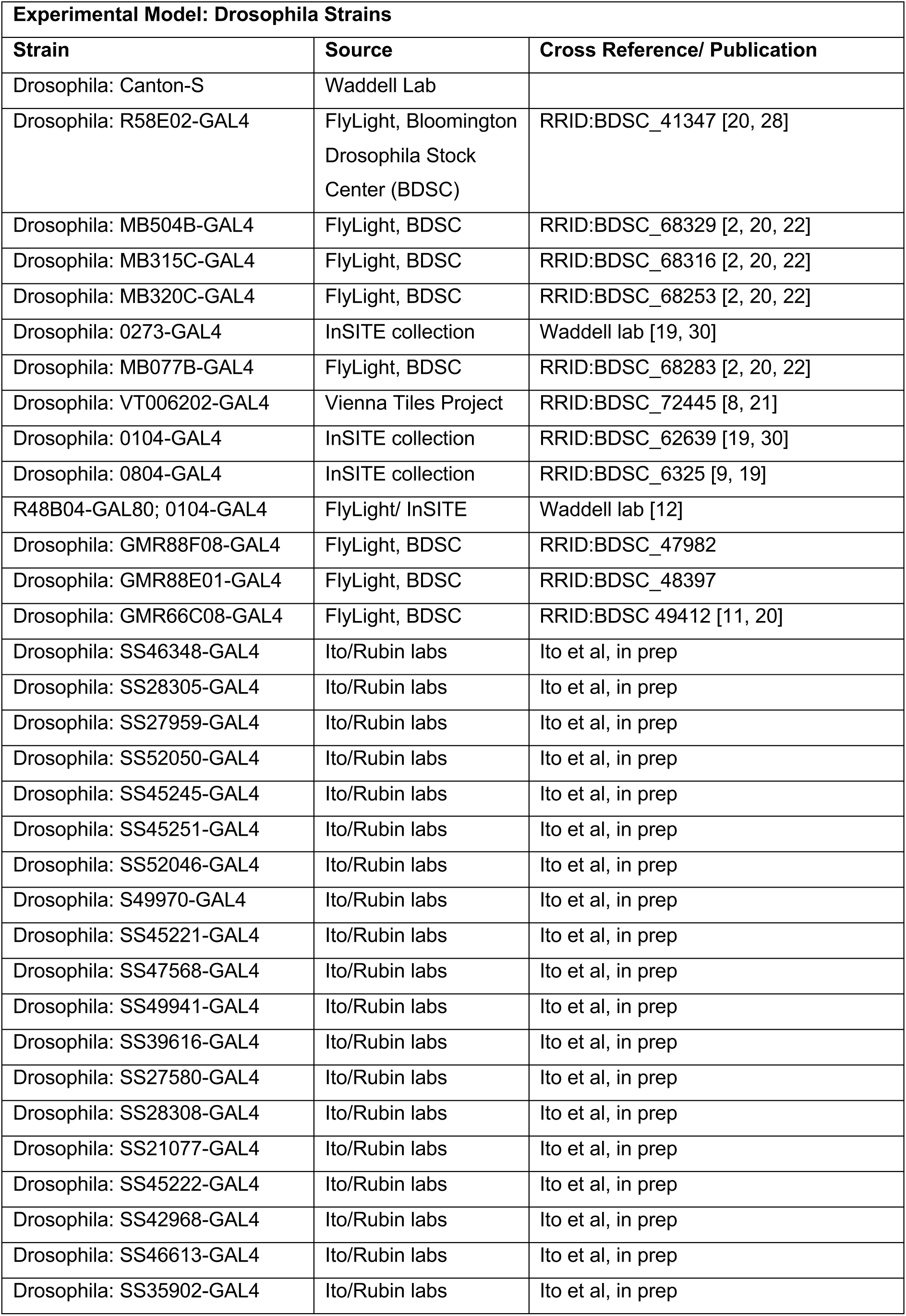
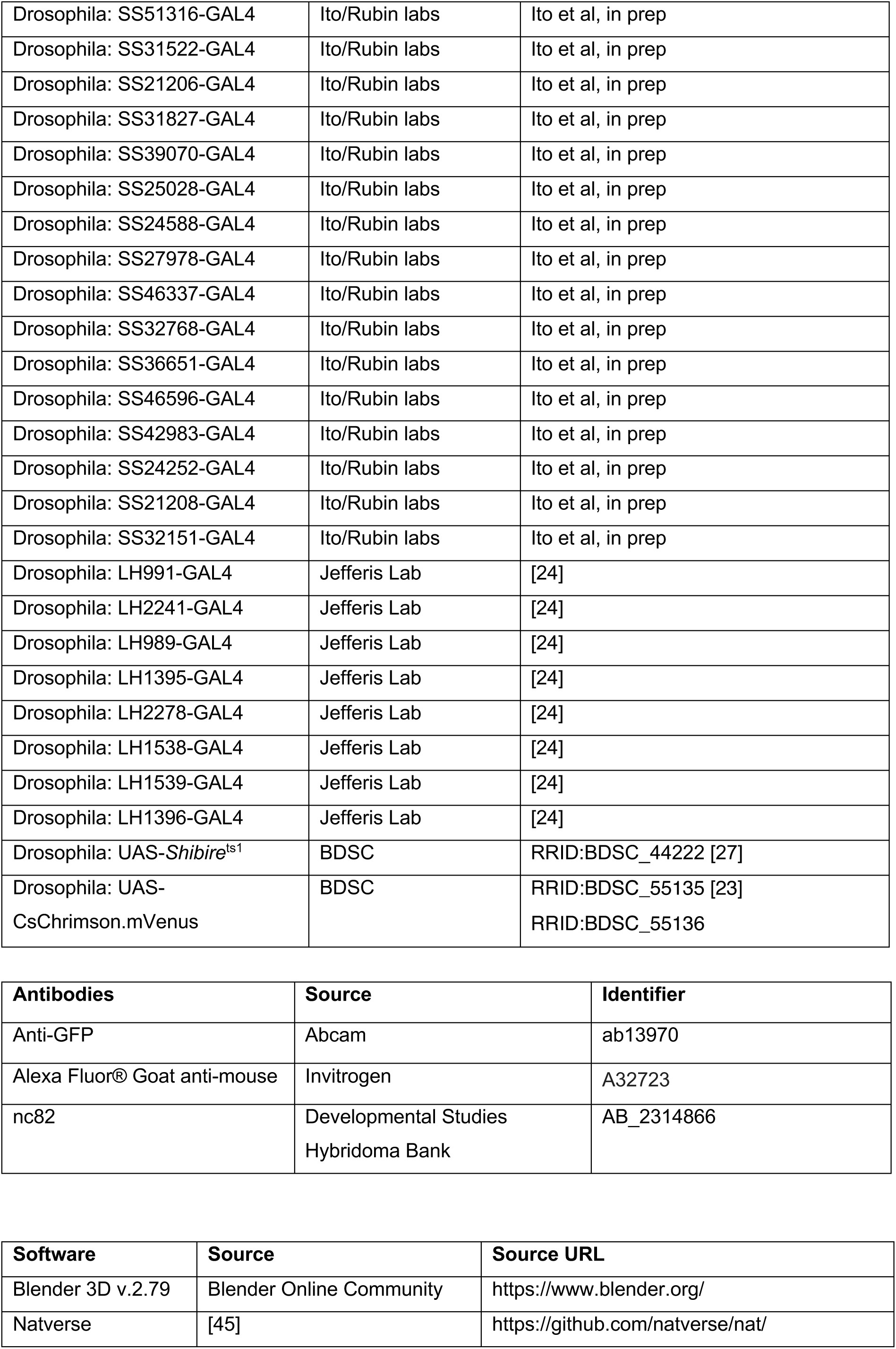
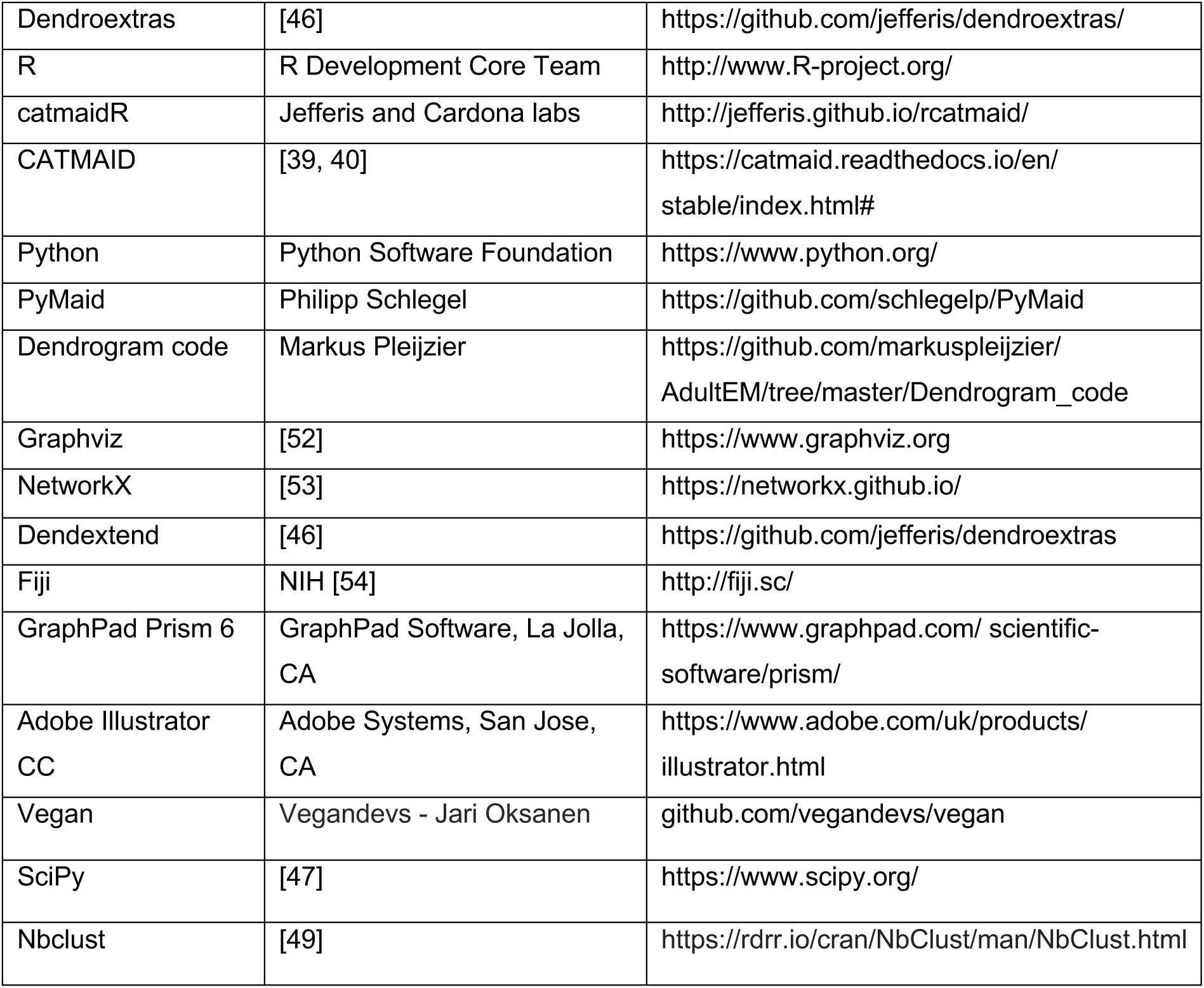

### METHOD DETAILS

#### Neuron reconstruction - ‘tracing’ in FAFB

Neurons were reconstructed from a serial section transmission electron microscopy (ssTEM) volume of a full adult fly (*D. melanogaster*) brain (FAFB) [4] using CATMAID, a web-based software for collaborative neural circuit reconstruction from large image datasets (https://catmaid.readthedocs.io/en/stable/) [39, 40]. Consistent with previous studies [3,37,41], tracing followed the centerline of a neuron’s profiles through the dataset to reconstruct neurite morphology and annotate synaptic sites. We used an established and tested iterative approach [40] where initial reconstruction is followed by a systematic proofreading by at least two expert reviewers (>500 h of tracing experience). We also took advantage of recent automatic segmentation efforts of the FAFB dataset [42], where flood-filling algorithms create volumetric segmentations of the EM data. These segmentations are then skeletonised to produce neuron fragments that can be joined together to expedite reconstruction. Human proof-reading is still required to remove incorrect merges of skeletons. In this study auto-segmentation was only used to aid the tracing of DAN input neurons to identification (see below).

Synaptic sites were identified based on three, previously described criteria [43] and reviewed as above: an active zone with (1) T-bar(s) and (2) surrounding vesicle cloud, and (3) a synaptic cleft to which all postsynaptic neurons must have access. In *Drosophila*, presynapses have been found on fine axonal processes [40], boutons [44], and other neurites that are neither in the dendritic nor the axonal field. Post-synapses have been found on large or fine dendritic processes and spine-like twigs that are shorter than 3 µm [40]. It has been estimated that the tracing approach employed finds 99.8% of presynapse and 91.7% of postsynapses [40]. The probability of identifying false-positive postsynapses is 2.2% and negligible for presynapses.

#### DAN identifying, tracing and quality control

DANs were first identified by selecting potential profiles in the midline commissure between left and right hemisphere MBs. These profiles were traced until axonal branches could be identified in the MB compartment of interest. We exhausted all possible profiles between the two MB compartments and in doing identified, both PPL1-γ1pedc DANs, and right hand side (RHS) β’2a DANs, and γ5 DANs. Although the number of γ5 DANs has been estimated to be between 8 and 21 (Aso et al., 2014) we nevertheless considered the possible existence of unilateral γ5 DANs, which would not have a process extending across the midline. To do this we also sampled neural profiles in the descending tract where the processes of γ5 DANs enter the MB lobe. However, we did not identify additional γ5 DANs.

Following identification DANs were traced and reviewed (as described above). Full details of tracing and review are provided in the Revision Status Table below. In brief, the RHS PPL1-γ1pedc DAN was completed and subjected to standard expert review. The γ1pedc dendritic field was further extensively reviewed. 20 PAM-γ5 DANs on the RHS and 9 on the left were reconstructed and the RHS neurons received standard expert review strategy as described above. 7 PAM-γ5 DANs received further extensive reviewed of their dendritic field. Of the 4 PAM-β’2 DANs 2 underwent both standard and additional extensive review, one only received standard review and a fourth was only partially reviewed. Any neurons that were not reviewed to this standard were excluded from the analyses. We note that it was more challenging to reconstruct DANs than many other neurons in the *Drosophila* brain. DAN dendrites are very thin and have a dark/granular texture, which increases the likelihood of missing branches and synapses. We therefore scrutinized completion and postsynapse annotation for 7 PAM-γ5 DANs (representing all morphological clusters), 2 PAM-β’2a DANs and the PPL1-γ1pedc DAN. Following this extended reconstruction and revision effort, we are confident that we have annotated all identifiable postsynapses on these selected DANs. Comparing data obtained from the regular review protocol to that from our extended review effort showed that regular review captured ∼30% of the postsynapses on more than 60% of all cable. We also analysed the placement of old (regular review) and new (added following extensive review) synapses, by measuring their geodesic (along-the-arbor) distance to the dendritic root (Figure S2H). This analysis showed that each round of additional review adds new synapses that are distributed along the arbor.

Lastly, we assessed whether uneven tracing of input connectivity altered the clustering of DANs by randomly downsampling (see below) the 9 extensively reviewed neurons to a level of inputs traced for the other regularly reviewed neurons. DANs could be similarly clustered following the downsampling, demonstrating that our DAN clustering results are unlikely to vary greatly with additional tracing of more input neurons.

#### Tracing neurons providing inputs to DANs

When a postsynapse was annotated on a DAN, a single-(seminal) node profile was placed in the centre of the presynaptic cell, unless a neuron or fragment was already present. To reconstruct upstream neurons from these seminal nodes we randomised the sampling order from each postsynapse within the total population on a neuron-by-neuron basis. For the reviewed PAM DANs (18 γ5 and 4 β’2a) we typically traced over 85% of the input neurons to identification from annotated postsynapses on a DAN arbor (see Revision Status Table). For the PPL1-γ1pedc DAN, we traced from 50% of annotated DAN postsynapses to identify the input neurons. Tracing inputs to this collection of γ5, β’2a and γ1pedc DAN postsynapses recovered 821 upstream neurons, some of which connect to multiple DANs in the traced groups. The tracing of the upstream neurons also varies in level of completeness but all neurons were traced to identify their microtubule containing backbone and were followed to a soma to retrieve their gross morphology.

#### Revision Status Table

Table details the review status of each DAN traced and analysed in this study. The total number of identified dendritic postsynapses on each DAN and the percentage of the postsynapses that are connected to an input neuron that was traced to identification are listed. Only DANs that were expert reviewed and have more than 50 identified input neurons were included in the analyses.

| Publication name | CATMAID Skeleton ID | Review status | Total dendritic post synapses | % of dendritic post synapses connected to input neuron | Included for analyses (total (inputs IDed) >50) |
| --- | --- | --- | --- | --- | --- |
| **γ5-1(fb)** | 3639761 | standard expert review<br>+ extensive review of the dendrite | 288 | 94.1 | ✓ |
| **γ5-5(fb)** | 3643424 | standard expert review<br>+ extensive review of the dendrite | 183 | 91.3 | ✓ |
| **γ5-6(fb)** | 5234721 | standard expert review<br>+ extensive review of the dendrite | 217 | 91.2 | ✓ |
| **γ5-19(uc)** | 6181246 | standard expert review<br>+ extensive review of the dendrite | 204 | 91.2 | ✓ |
| **γ5-20(uc)** | 3134936 | standard expert review<br>+ extensive review of the dendrite | 322 | 89.4 | ✓ |
| **γ5-23(uc)** | 11262395 | standard expert review<br>+ extensive review of the dendrite | 186 | 88.2 | ✓ |
| **γ5-24(uc)** | 11274761 | standard expert review | 148 | 82 | ✓ |
| **γ5-25(uc)** | 11149656 | standard expert review (4%) | 137 | 90 | ✓ |
| **γ5-16(lc)** | 6019056 | standard expert review | 268 | 45 | ✓ |
| **γ5-10(lc)** | 5652208 | standard expert review | 247 | 81 | ✓ |
| **γ5-12(lc)** | 5914197 | standard expert review | 225 | 52 | ✓ |
| **γ5-2(lc)** | 3025479 | standard expert review | 123 | 87 | ✓ |
| **γ5-7(lc)** | 3026119 | standard expert review | 106 | 96 | ✓ |
| **γ5-4(lc)** | 3143746 | standard expert review | 95 | 58 | ✓ |
| **γ5-13(lc)** | 5952626 | standard expert review | 72 | 97 | ✓ |
| **γ5-21(da)** | 11169413 | standard expert review (60%) | 109 | 77 | ✓ |
| **γ5-15(ba)** | 6013704 | standard expert review | 145 | 93 | ✓ |
| **γ5-17(ba)** | 6059582 | standard expert review<br>+ extensive review of the dendrite | 239 | 94.1 | ✓ |
| **β'2a-1(2)** | 7777207 | standard expert review<br>+ extensive review of the dendrite | 307 | 92.5 | ✓ |
| **β'2a-2(nc)** | 7270967 | standard expert review<br>+ extensive review of the dendrite | 169 | 91.1 | ✓ |
| $\beta'2a-4(1)$ | 8317886 | standard expert review | 281 | 93 | ✓ |
| $\beta'2a-3(2)$ | 7512499 | standard expert review (30%) | 212 | 45 | ✓ |
| **PPL1-γ1pedc** | 1159799 | standard expert review<br>+ extensive review of the dendrite | 3480 | 46.9 | ✓ |
| γ5-22 | 11175371 | standard expert review | 94 | 46 | |
| γ5-8 | 5631471 | standard expert review (73%) | 44 | 59 | |
| $\beta'2a-5$ | 6969526 | not reviewed (not fully<br>reconstructed) | 3 | 33 | |
| $\beta'2a-6$ | 5886218 | not reviewed (not fully<br>reconstructed) | 4 | 75 | |

#### Analysis of neuroanatomical features and DAN connectivity

All analyses were performed in R and Python using open-source software. PyMaid (https://github.com/schlegelp/PyMaid) and RCatmaid (http://jefferis.github.io/rcatmaid/; http://jefferis.github.io/elmr/) were used to interface with CATMAID servers and perform morphological analyses. Neuron analyses were performed with natverse functionality (https://github.com/natverse/nat) [45] or custom-written code, which is available on request. Hierarchical clustering was performed using base R functions and dendrogram visualisations with dendroextras (https://github.com/jefferis/dendroextras). Tanglegrams were generated using dendextend (https://github.com/talgalili/dendextend) [46]. Mantel test analyses were computed with the vegan R package (https://github.com/vegandevs/vegan).

#### Clustering

##### Euclidean and Manhattan distance metrics for clustering

To analyse and draw conclusions from differences and similarities in large amounts of connectivity or morphology data, the information is represented in the form of distance matrices between each data point in space. Euclidean distance is the direct (bee-line) distance between two points in a Cartesian coordinate system. Manhattan distance between two data points in a Cartesian system is the sum of distances between the coordinates.

##### Ward’s clustering criterion

Ward’s method was used for agglomerative hierarchical clustering (part of R base package). Each datapoint starts in its own cluster and pairs of clusters are merged, moving up the hierarchy. At each step the pair of clusters with minimum within-cluster variance are merged. Connectivity data as well as morphology data was clustered using Ward’s criterion to compare to clusters formed using average linkage criterion.

##### Average linkage clustering criterion

Average linkage (also known as unweighted pair group method with arithmetic mean, UPGMA) is another criterion for agglomerative hierarchical clustering. With average linkage clustering pairwise dissimilarities between each element in cluster 1 and 2 are computed and the average of these dissimilarities are considered as the distance between the two clusters. Clusters separated by the smallest distance are merged during clustering, moving up the hierarchy. Both morphology and connectivity data were clustered using average linkage to compare to data clustered with Ward’s criterion.

##### Clustering DANs by morphology

To compensate for different levels of completeness of tracing, DANs were simplified to their longest tree with 200 branch points (the minimum number of branch points throughout the PAM DAN population). Morphological similarity matrices were calculated using NBLAST [18]. Hierarchical clustering was primarily performed using base R functions, taking Euclidean distance matrices of similarity scoring, with average linkage clustering criterion. Morphology clustering was performed with Ward’s and average linkage criteria for comparison.

##### Clustering DANs by input connectivity

Connectivity information was retrieved from CATMAID after synapse annotation and upstream tracing of input neurons. Only neurons upstream of the dendritic region of DANs with >50 sampled profiles were included in the analyses. Before clustering the number of synapses annotated on each DAN was normalized to reduce bias in clustering that could arise from the varying levels of tracing completeness and/or natural differences in the number of inputs to the different DANs. Hierarchical clustering was primarily performed using the Manhattan distance between upstream connectivity profiles of DANs with Ward’s clustering criterion. Connectivity data was also clustered using the average linkage criterion for comparison.

##### DAN PPL-γ1pedc postsynapse clustering

For PPL1-γ1pedc the x, y and z coordinates of postsynapses on the dendrite were clustered using Ward’s hierarchical clustering in the SciPy package (https://www.scipy.org/, [47]).

##### Silhouette method to determine accuracy of the number of clusters

Silhouette is a graphical representation of the quality of clustering across a range of potential values for k (the number of clusters). Where possible Silhouette was used to select an appropriate number of clusters (it was less useful for morphology clustering). This method measures how similar observations are to their own cluster and how dissimilar to other clusters. The average silhouette width ranges from 0 and 1, with 1 indicating observations are well clustered. To validate DAN clustering the average silhouette width was calculated using the nbclust R package [48, 49].

##### Input neuron morphology clustering

Morphology clustering of upstream neurons was performed using hierarchical clustering with average linkage criterion. This involved a multi-step approach to account for varying levels of tracing and for the morphological diversity of 821 neurons (see Figure below). Coarse clustering was performed taking the soma tract as the primary feature of neuron identity. Subsequently the larger primary clusters were subclustered by splitting neurons into the primary neurite and its complement/remainder. Similarity matrices were calculated using NBLAST and an element-wise mean (80:20) was used for clustering. For fine clusters, weighting methods were selected iteratively depending on overall sub-cluster morphology.

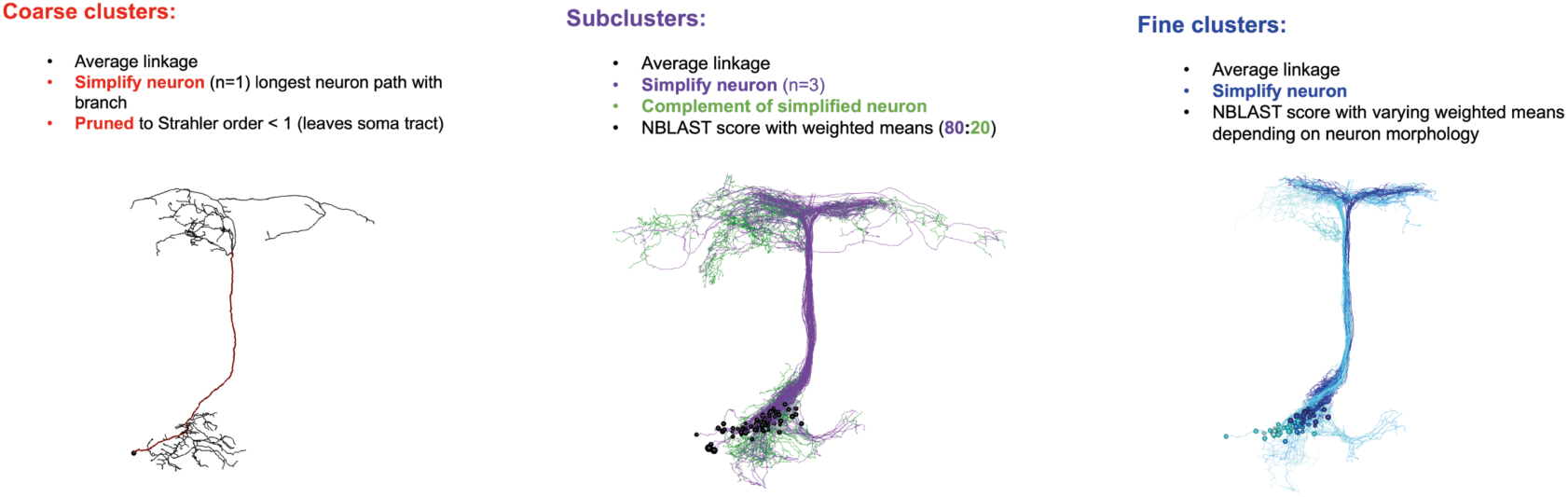
*A multistep clustering approach to classify DAN input neurons*. (Left) Coarse clusters are generated by hierarchical clustering of an ‘all-by-all’ NBLAST comparison of simplified neurons. Neurons are simplified leaving the soma tract and the main neurite. This first step works unsupervised on all 821 neurons, resulting in 20 coarse clusters (Figure 3I). Middle: Larger primary clusters are subclustered by weighting defining neural features. 80% towards the core neurons and 20% towards the complement of the core neuron - “the fine branches” (green). This example shows SEZONs which are comprised of long tracts and a simple branched arbor. This created 153 sub-clusters with little human supervision. (Right) Fine clustering. Neurons in each sub cluster have different key morphological features thus subsequent clustering steps are different in exact parameters. These settings are reviewed manually. We determined the step - fine clustering – in most cases to be sufficiently accurate to separate all neurons that are morphologically different in different fine clusters.

##### Tanglegrams to compare clustering of 2 feature spaces

Tanglegrams were generated to visually compare clustering dendrograms produced by different criteria (eg. average linkage versus Ward’s) or clustering based on morphology versus those produced using connectivity. Dendrogram layouts were determined to minimise edge crossing (i.e. minimise Manhattan distance between corresponding DANs) using dendextend [46].

##### Mantel test to determine dependence of 2 feature spaces

The Mantel test was used to compare 2 sample spaces - here neuron morphology distance matrices obtained from all-by-all NBLAST and distances based on connectivity were used. To create distance matrices for connectivity, connectivity matrices were normalized by the postsynaptic budget of DANs. The implementation of the Mantel test was based on [50]. Pearson’s correlation between the two observed datasets was calculated, then one of the matrices was shuffled 10^7^ times and each event tested for correlation with the observed data. The number of events where the correlation is higher than between the two original datasets was divided by the amount of comparisons (10^7^) to create a p-value. When p-values were lower than the significance level, it was concluded that the null model of independence between the two feature spaces could be rejected.

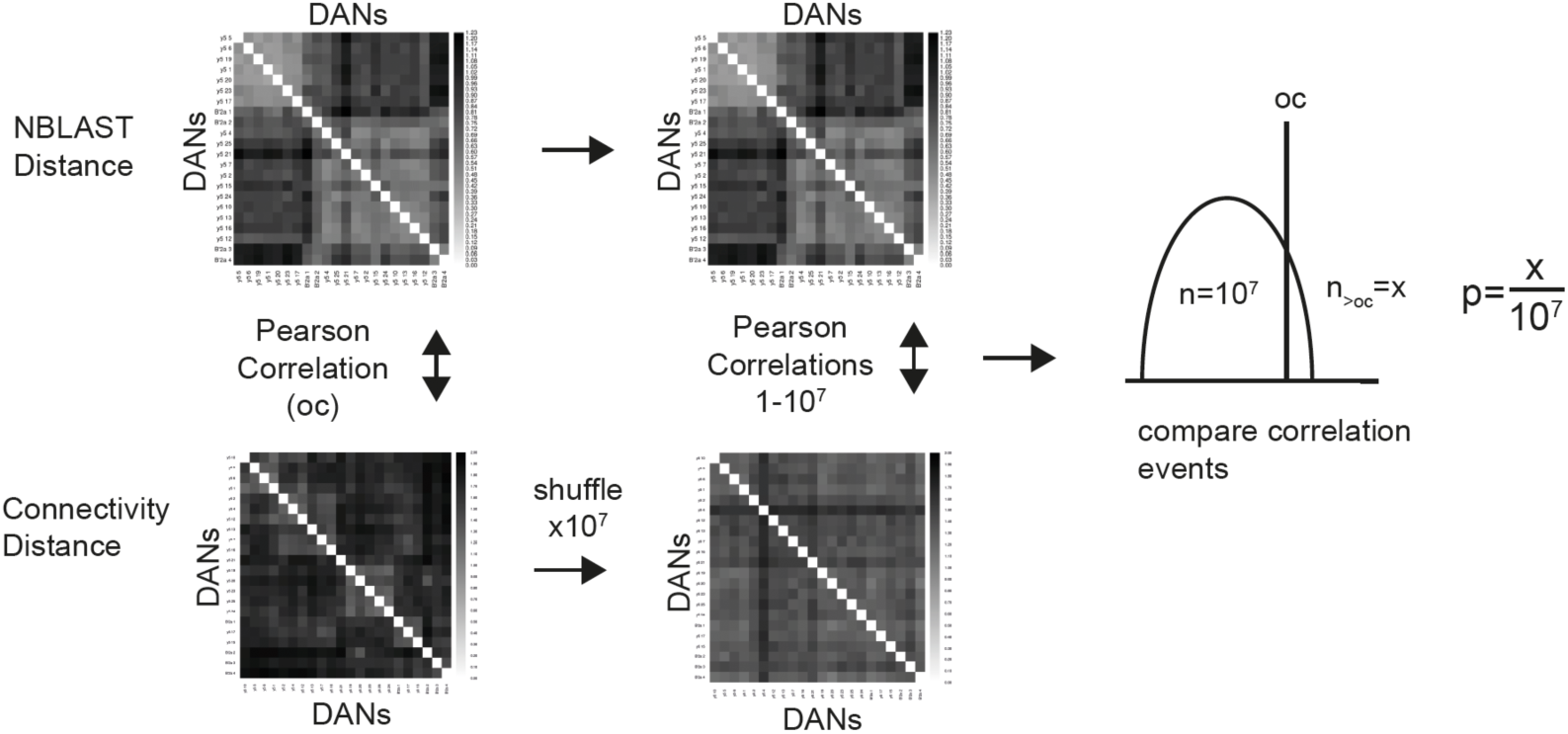
M*a*ntel *test to determine statistical significance of correlations between properties (cluster results) compared in tanglegrams.* Here the morphological NBLAST distance matrix is compared to the observed connectivity distance matrix in Figure 2E. Comparison is by Pearson’s correlation of the Manhattan distributions. The Connectivity matrix is subsequently shuffled 10^7^ times and the resulting matrices are compared to the observed NBLAST distance matrix. Lastly (right), the correlation coefficients that are higher than the correlation coefficient between the observed data (morphology and connectivity) are divided by the number of randomisations (10^7^) to obtain a p-value. A small p-value suggests the compared distance matrices are correlated.

#### Downsampling connectivity to verify reproducibility of clustering

To verify that clustering into the observed DAN groups does not result from bias in the relative completeness of tracing of the input network to each DAN, down-sampled the datasets so that all DANs were randomly stripped of 5-50% of their input connections. Clustering resulting from 10,000 repetitions of this down-sampling were then compared to the clustering obtained from the full dataset. In addition, to exclude that more exhaustive reviewing of a few exemplary DANs might skew the clustering, we created a dataset of 10,000 repetitions where the connectivity of only the exhaustively reviewed DANs was reduced to the average of all the remaining DANs. Clustering obtained with this normalised dataset was also compared to that retrieved using the full dataset.

#### Cluster similarity analyses

In 10,000 iterations of resampling a dataset with reduced connectivity, each neuron has a different likelihood to cluster with the same original group that it did in the 100% connectivity dataset or, with any of the other original groups. We therefore also calculated the average likelihood (over the 10,000 trials) that a downsampled DAN clustered with the same group that it clustered with in the full 100% connectivity dataset. These values were then plotted as a stacked bar plot.

#### Fowlkes-Mallows Index (FMI)

The Fowlkes-Mallows Index measures the similarity of the content between two different clusterings. The performance of the first clustering is compared to that of the second clustering (which is assumed to be perfect). Exact matches/good performance result in an FMI = 1 [51].

#### Neuropil of origin

To identify SEZONs and LH-associated neurons, the cable length within the respective neuropil mesh (3D bounding box) was calculated. A cut off of >60 nodes within the defined neuropil region was required for classification.

#### Dendrogram representations

Dendrogram representations of neurons were created as in [11]. Dendrograms are 2D representations of 3D neuronal reconstructions which preserve the topology of neuron and visualise specific synapses on specific branches. The neato layout (Graphviz, https://graphviz.gitlab.io/ [52]) attempts to minimize a global energy function, equivalent to statistical multi-dimensional scaling to represent the neuron morphology as a graph. Code available (https://github.com/markuspleijzier/AdultEM/tree/master/Dendrogram_code) using the Graphviz library with Python bindings provided by NetworkX, (https://networkx.github.io/ [53]).

#### Marking MBON postsynapses on dendrograms by closest DAN cluster

Euclidean distances between MBON-γ5β’2a postsynapses and the closest DAN presynapse were measured and marked with the identity of the morphological DAN clusters (Figure 1). The Euclidean distances were then thresholded to within 2 µm and the synapses identified to be under that threshold were plotted on a neato dendrogram. The plot in Figure S4F therefore shows all postsynapses within a 2 µm diffusion distance from a dopaminergic presynapse.

#### Edge weight distribution

Edge weight distributions describe how many upstream neurons contribute a given number of presynapses to a connection with a postsynaptic neuron (frequency vs number of synapses). Normalising by the total number of postsynapses details the percentage of the total postsynaptic budget a given number of synapses represents. For example, if a neuron makes 10 presynapses onto a postsynaptic neuron, which has a total of 100 postsynapses, then that upstream neuron contributes 10% of the postsynaptic budget.

#### DAN-MBON direct connectivity

Identified MBONs were collapsed by type. The number of synapses between DANs and MBONs was normalized by the number of all DAN-MBON connections of the given DAN. Connectivity matrices can be calculated for single branches of a neuron after defining the relevant branchpoints in CATMAID. For the PPL1-γ1pedc DAN, we manually split the dendrite into 4 postsynaptic clusters, as defined from cluster analyses, and recorded the specific connectivity to each of these clusters/branches.

#### DAN connectivity similarity matrices

A connectivity similarity score between 2 DANs was defined as one minus half of the Manhattan distance between their normalised connectivity patterns (normalised connectivity patterns of DANs shown in Figure 2E).

#### Statistical analysis of DAN connectivity - comparison to a null model of random connectivity

A DAN input connectivity matrix was first randomised 10^4^ times, respecting both DAN postsynaptic budget and input neuron presynaptic budget, so that after randomization each row sum and each column sum remained the same as in the observed data (i.e. each DAN gets the same number of inputs and each input neuron has the same number of outputs). Then the Manhattan distance between upstream connectivity profiles of DANs in the observed data and those in simulated random matrices, both normalised by DAN postsynaptic budget were calculated and means of these distances were compared to obtain a p-value describing the similarity of these means. A p-value lower than the significance level concluded that the null model of randomized connectivity could be rejected.

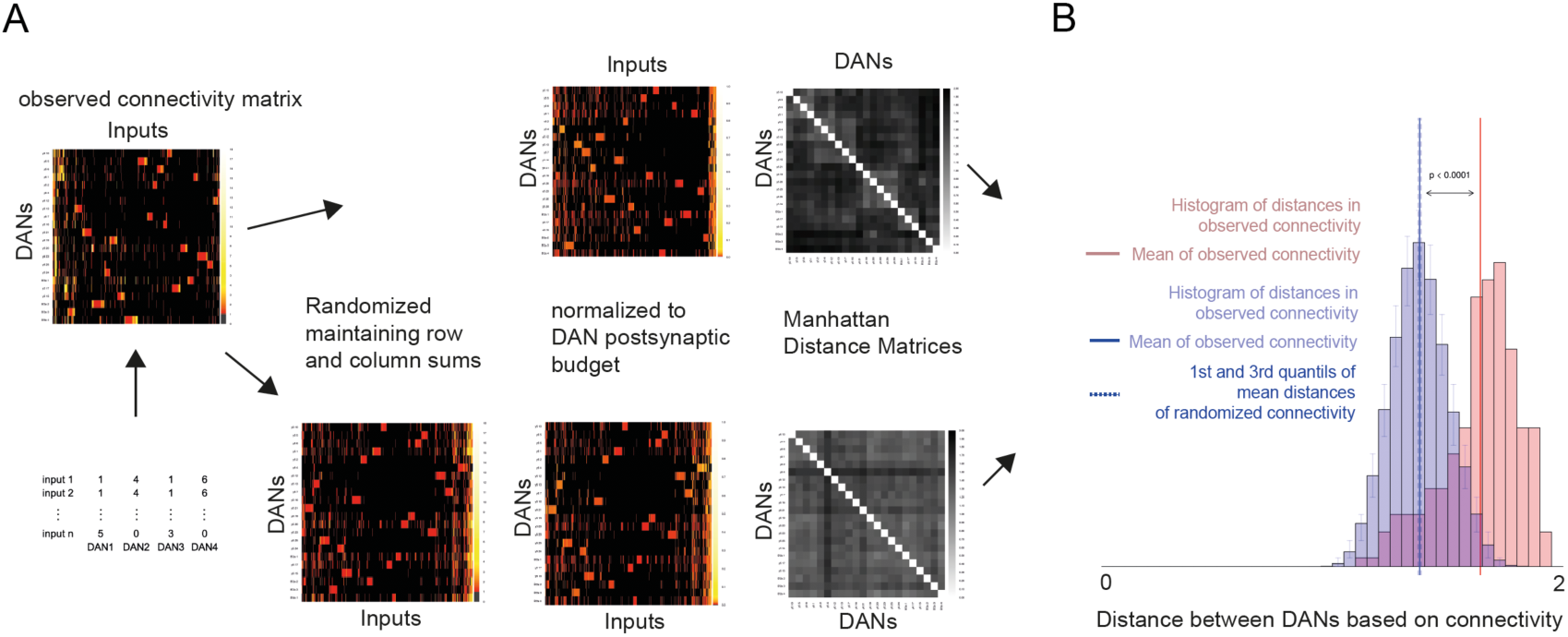
R*a*ndom *connectivity null model*: CATMAID synaptic connectivity from observed data is represented as a connectivity matrix (left). This matrix is randomised conserving the proportion of DAN postsynaptic budget and input presynaptic budget in the network. 10^4^ random matrices are computed for both the observed connectivity matrix and the random matrix, normalised by DAN postsynaptic budget (middle). A Manhattan distances between observed and simulated matrices are calculated. (B) Statistical analysis of mean Manhattan distances is carried out by comparing the distribution of Manhattan distances in the observed data (red) with the mean distribution of 10^4^ simulated matrices (blue). The means of the two distributions are significantly different; p<0.0001. Therefore, the organization of DAN input connectivity is not random.

#### Data visualization

Reconstructed neurons were visualized using Blender 3D, an open-source 3D software (https://www.blender.org/) or natverse [45] and RCatmaid packages in R (http://jefferis.github.io/rcatmaid/). Neuron data from CATMAID were imported using an existing CATMAID plugin for Blender (https://github.com/ schlegelp/CATMAID-to-Blender [41]). Maximum intensity projections of Confocal stacks were generated using FIJI [54].

#### 3D Representations and Videos

3D representations of traced skeletons that were obtained as swc files from CATMAID after reconstruction were either plotted using the natverse R tool box, or 3D representations were created and rendered with blender v2.79. Video footage of skeletons was rendered to obtain 3D representations with blender v2.79 and cut with Adobe Premiere Pro2020.

#### Matching genetic driver lines to EM skeletons

EM skeletons were matched to published library GAL4 lines using registered Janelia FlyLight micrograph data from VirtualFlyBrain (https://v2.virtualflybrain.org). NBLAST similarity matrices were calculated comparing both data types and top hits were visualised and manually cross compared. In case of new split-GAL4 SS lines created by Masayoshi Ito (Ito et al., in prep), imaging stacks were newly registered with bridging and mirror registrations from the natverse package [45].

#### Immunohistochemistry and imaging

Following [9], Brains were dissected in cold Phosphate Buffered Saline (PBS) and fixed in PBS with 4% paraformaldehyde at 25 °C for 40 min. They were washed 3X by quick PBS exchange and 3X for 20 min each, in PBS containing 0.5% Triton-X100 (PBT), followed by 30 min incubation in PBT containing 5% normal goat serum.

Brains were either then imaged for endogenous expression of GFP or the buffer was exchanged with anti-GFP (1:200; Abcam) and anti-nc82 (1:50; DSHB) antibodies in PBS containing 0.5% Triton-X100 (PBT). Brains were incubated for 24-72 h at 4 °C, washed 3X 20 min in PBT at 25°C, followed by incubation in PBT containing the appropriate Alexa Fluor® secondary antibodies (Invitrogen) overnight at 4 °C. Brains were then washed 3X 20 min in PBT at 25°C, before being mounted on slides with VectaShield® (Vector Labs). Imaging was performed using a Leica TCS SP5X confocal microscope.

#### Behavioural experiments

All *Drosophila* strains were raised at 25°C on standard cornmeal-agar food at 50-60% relative humidity in a 12:12 h light:dark cycle. For experiments, female flies bearing effector transgenes were crossed to male flies bearing GAL4 drivers. For driver line details see Key Resources Table. Driver lines were obtained from the Janelia FlyLight collection [20] or split-GAL4 [55] collections, the Vienna Tiles Project [21], and the InSITE collection [19]. New split-GAL4 ‘SS’ lines were created in the Ito/Rubin labs (Ito et al., in prep). UAS-*CsChrimson* [23] and *UAS-Shibire^ts^* [27] effectors were used to stimulate and block specific neurons. For wildtype controls Canton-S flies were used. For heterozygous controls, GAL4 lines were crossed to Canton-S flies.

The odors used for US substitution, sucrose learning and DEET learning experiments were 10^-3^ dilutions of 3-octanol (OCT) and 4-methylcyclohexanol (MCH) in mineral oil. For extinction experiments odor concentrations of 10^-6^ were used to avoid pre-exposure effects [11]. Experiments were performed at 23°C and 55-65% relative humidity, except for electric shock learning which occurred at 70% relative humidity.

#### US-substitution experiments using CsChrimson

In both the behavioural screen and follow-up experiments, neurons were artificially activated to substitute for an unconditioned stimulus in the training chamber of a T-maze. Prior to the experiments, 80-120 1-5 day old mixed sex flies were housed on standard food supplemented with 1% all-trans-Retinal for 3 days before a 20 – 28 h starvation period in vials containing 2 ml 1% agar as a water source and a 2×4 cm strip of Whatman filter paper. During training, groups of flies were exposed to the CS- for 2 min followed by 30 s rest with fresh air, then 2 min of CS+ odor with optogenetic activation of the genetically encoded Channel Rhodopsin with red light exposure.

Three red (620-630nm) LEDs (Multicomp, p/n OSW-4338) with 3 W maximum power were mounted on the training arm of a T-maze and 1ms pulses were driven at 1.2V with a stimulation frequency of 500Hz, which is flicker free red-light that flies cannot see. For screening, immediate memory testing followed. Flies were transferred back into their starvation vials after training before testing 30 min memory.

#### Appetitive olfactory learning with sucrose reward

Prior to the experiments, 80-120 3-8 day old mixed sex flies were starved for 20-28 h in vials containing 2 ml 1% agar as a water source and a 2×4 cm strip of Whatman filter. Flies were transferred to 32 °C 30 min before training. During training, groups of flies were exposed to the CS- odor with dry paper for 2 min followed by 30 s of fresh air, then 2 min of CS+ odor exposure with dry sugar paper. Flies were either tested immediately after training or were transferred back into 25°C starvation vials after training prior to testing 30 min memory.

#### Aversive olfactory learning with bitter reinforcement

Flies were aversively trained with DEET as previously described [16]. In brief, prior 80-120 3-7 day old mixed sex flies were starved for 20-24 h in vials containing 2 ml 1% agar and a 2×4 cm strip of Whatman filter paper. Training and immediate testing were performed at 32°C. During training groups of flies were exposed to the CS- odor with 1% agar on filter paper for 2 min followed by 30 s fresh air, then 2 min of CS+ odor with 0.4% DEET, 3 M xylose and 100 mM sucrose in 1% agar on filter paper.

Flies were tested for their odor preference immediately after training.

#### Aversive memory extinction

Extinction memory was tested as described [11]. In brief, mixed sex groups of 80-120 flies were transferred into vials with 2 ml cornmeal medium and a 2×4 cm strip of Whatman paper for 18-26 h before training. Aversive olfactory conditioning in the T-maze was conducted as previously described [31]. Flies were exposed to the CS+ odor for 1 min paired with twelve 90 V electric shocks at 5 s intervals. Following 45 s of clean air, the CS- odor was presented for 1 min without shock. Immediately after training flies were transferred to 32°C. 30 min later flies were re-exposed twice to either the CS- or CS+ odor with a 15 min interval. Flies were then returned to permissive 23°C and tested 15 min later for memory performance.

#### Memory testing and statistical analyses of behavioural data

To test memory performance flies were loaded into the T-maze and transported to the choice point where they were given two min to choose between the CS+ and CS- odors in the dark. A Performance Index was calculated as the number of flies in the CS+ arm minus the number in the CS- arm, divided by the total number of flies [31]. MCH and OCT, were alternately used as CS+ or CS- and a single sample, or n, represents the average performance score from two reciprocally trained groups. Statistical analysis was carried out with GraphPad, Prism (v8.1). All experiments were analysed with a one-way ANOVA. For extinction CS- re-exposure flies were compared with CS+ re-exposure flies and Tukey’s post-doc analyses for multiple comparisons applied. For all other experiments statistical comparisons were performed between a wildtype control and Dunnett’s post hoc analysis carried out for multiple comparisons.

