## Supplemental figures for "Input connectivity reveals additional heterogeneity of dopaminergic reinforcement in *Drosophila*"

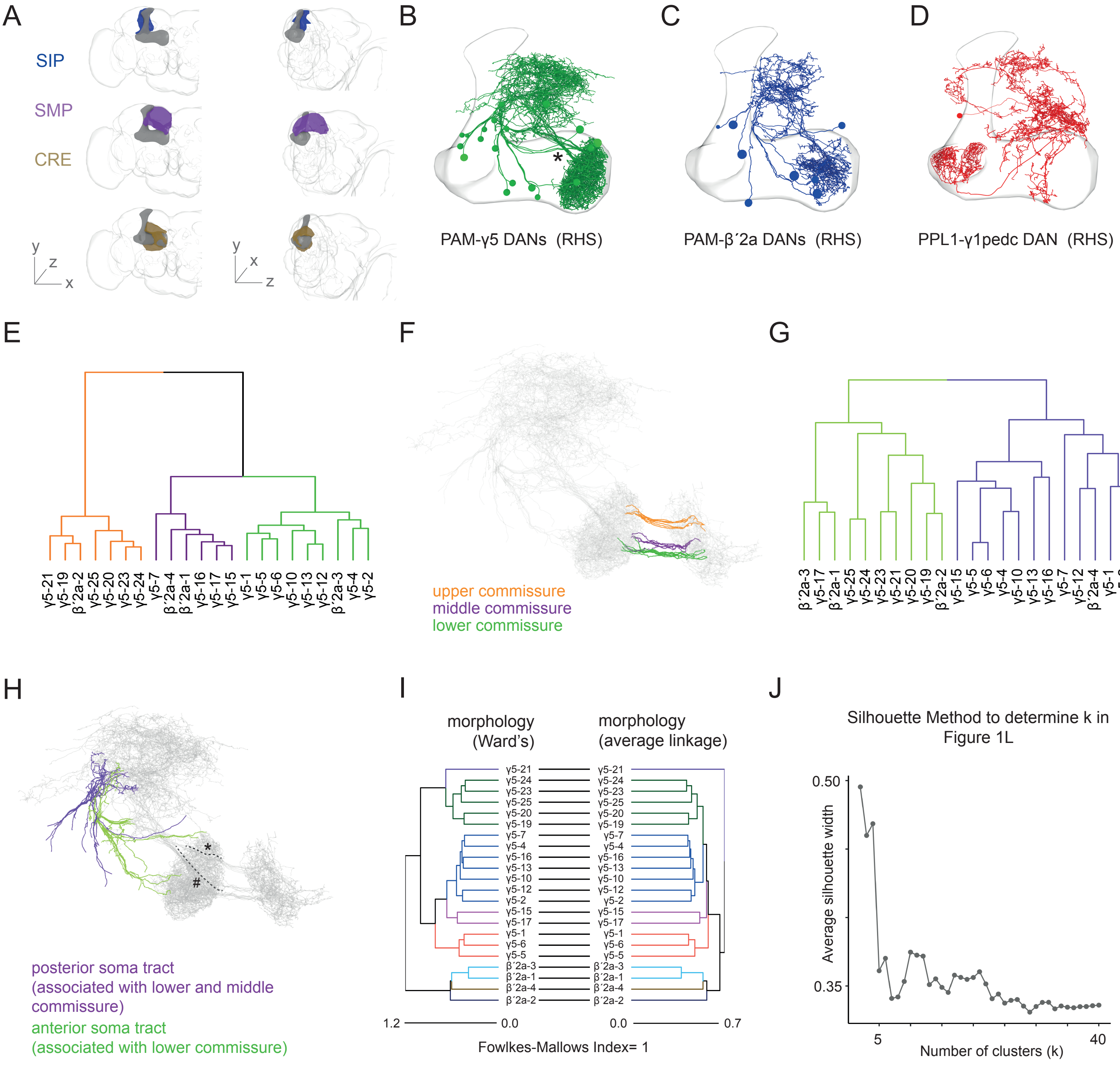

A
SIP
SMP
CRE
y z x
y x z
B
PAM-γ5 DANs (RHS)
C
PAM-β´2a DANs (RHS)
D
PPL1-γ1pedc DAN (RHS)
E
γ5-21 γ5-19 β´2a-2 γ5-25 γ5-20 γ5-23 γ5-24 γ5-7 β´2a-4 β´2a-1 γ5-16 γ5-17 γ5-15 γ5-1 γ5-5 γ5-6 γ5-10 γ5-13 γ5-12 β´2a-3 γ5-4 γ5-2
F
upper commissure
middle commissure
lower commissure
G
β´2a-3 γ5-17 β´2a-1 γ5-25 γ5-24 γ5-23 γ5-21 γ5-20 γ5-19 β´2a-2 γ5-15 γ5-5 γ5-6 γ5-4 γ5-10 γ5-13 γ5-16 γ5-7 γ5-12 β´2a-4 γ5-1 γ5-2
H
posterior soma tract
(associated with lower and middle commissure)
anterior soma tract
(associated with lower commissure)
I
morphology (Ward's)
morphology (average linkage)
γ5-21 γ5-24 γ5-23 γ5-25 γ5-20 γ5-19 γ5-7 γ5-4 γ5-16 γ5-13 γ5-10 γ5-12 γ5-2 γ5-15 γ5-17 γ5-1 γ5-6 γ5-5 β´2a-3 β´2a-1 β´2a-4 β´2a-2
1.2
0.0
0.0
0.7
Fowlkes-Mallows Index= 1
J
Silhouette Method to determine k in Figure 1L
Average silhouette width
0.50
0.35
5
40
Number of clusters (k)

**Figure S1 related to Figure 1**

(A) Areas of neuropil providing most input to the DANs in this study as defined in CATMAID and named according to [17]. The superior medial protocerebrum (SMP, purple), superior intermediate protocerebrum (SIP, blue) and crepine (CRE, ochre) are shown from the front (left) and form the midline (right). The MB and whole brain neuropil are depicted in grey. (B-D) Representations of all reviewed DANs in the fly's right hemisphere. (B) 20 PAM-γ5 DANs. MB outlined, asterisk indicates 2 descending soma tracts. (C) 8 confirmed PAM-β´2a DAN candidates. (D) PPL1- γ1pedc DAN with elaborate dendrite outside the MB. (E) Dendrogram using only the commissure part of the PAM-γ5 and PAM-β´2a DANs reveals 3 morphological clusters. (F) Projection view of the upper (orange), middle (purple) and lower (green) commissure DAN groups from the clustering in (E).Full DAN morphologies are shown in grey. (G) Average linkage clustering using the tracts connecting the somata and dendrites reveals 2 distinct groups. (H) Projection view of the posterior (purple) and anterior (green) soma tracts from clustering in (G). The posterior tract mostly contains lower commissure DANs (dashed line and hash) whereas the anterior tract mostly houses upper commissure DANs (dashed line and asterisk). (I) Tanglegram illustrating comparison of morphological clustering obtained using average linkage criterion (as in Figure 1B) to that retrieved using Ward's criterion. The clusters are identical, hence Fowlkes-Mallows Index is 1. Note: the between cluster relationship varies slightly. (J) Silhouette method to determine the appropriate number of clusters (k) selected for Ward's criterion based hierarchical clustering of 3-dimensional positions of PPL1-γ1pedc post synapses. Accuracy drops after 4 clusters and k=4 best reflects the quadripartite structure of the PPL1-γ1pedc dendrite.

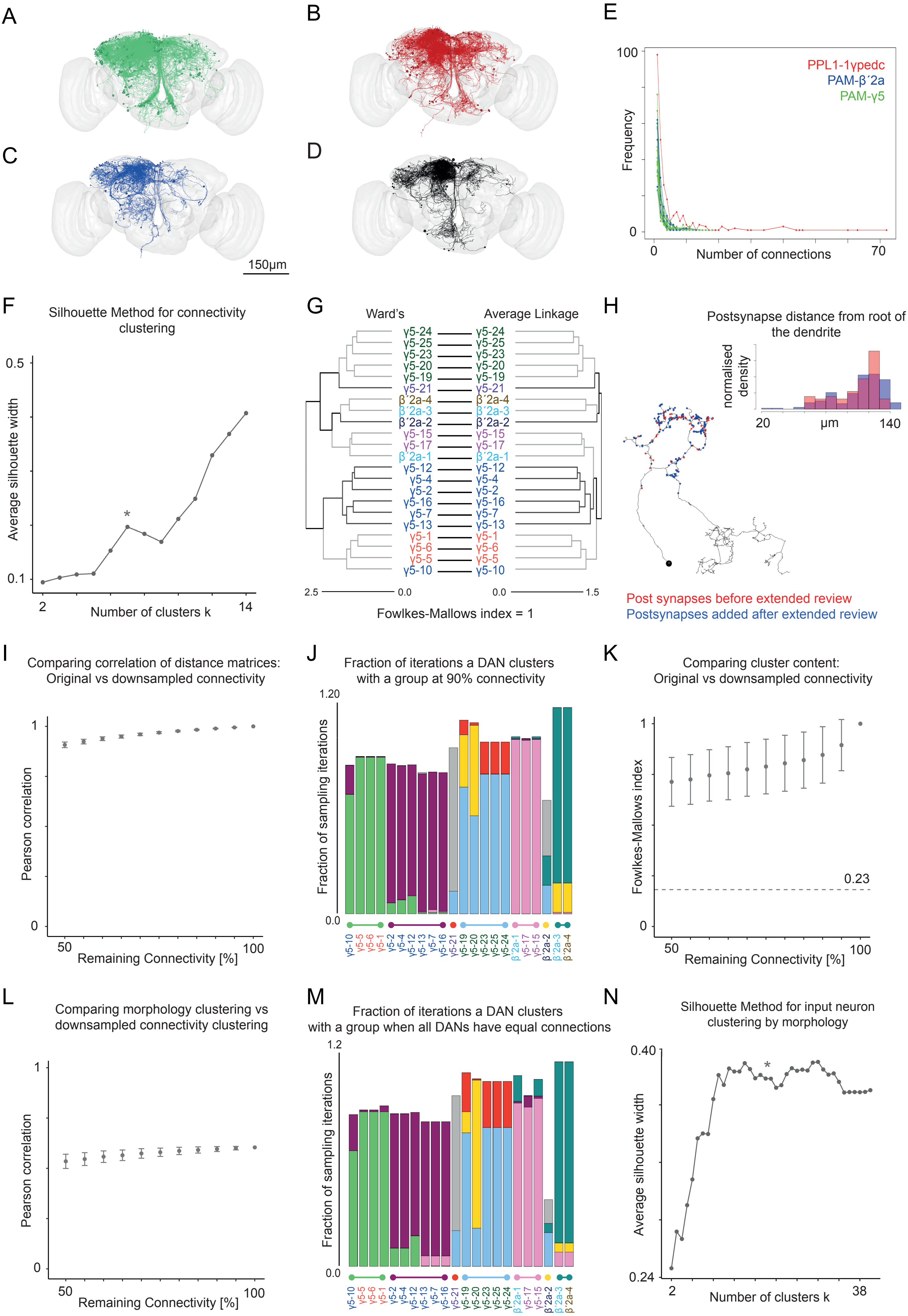
A
B
C
D
E
150µm
100
0
Frequency
Number of connections
70
PPL1-1γpedc
PAM-β´2a
PAM-γ5
F
Silhouette Method for connectivity clustering
Average silhouette width
0.5
0.1
*
2
Number of clusters k
14
G
Ward's
Average Linkage
γ5-24
γ5-25
γ5-23
γ5-20
γ5-19
γ5-21
β´2a-4
β´2a-3
β´2a-2
γ5-15
γ5-17
β´2a-1
γ5-12
γ5-4
γ5-2
γ5-16
γ5-7
γ5-13
γ5-1
γ5-6
γ5-5
γ5-10
2.5
0.0
0.0
1.5
Fowlkes-Mallows index = 1
H
Postsynapse distance from root of the dendrite
normalised density
20
µm
140
Post synapses before extended review
Postsynapses added after extended review
I
Comparing correlation of distance matrices: Original vs downsampled connectivity
Pearson correlation
1
0
50
Remaining Connectivity [%]
100
J
Fraction of iterations a DAN clusters with a group at 90% connectivity
1.20
Fraction of sampling iterations
0.0
γ5-10 γ5-5 γ5-6 γ5-1 γ5-2 γ5-4 γ5-12 γ5-13 γ5-7 γ5-16 γ5-21 γ5-19 γ5-20 γ5-23 γ5-25 γ5-24 β´2a-1 γ5-17 γ5-15 β´2a-2 β´2a-3 β´2a-4
K
Comparing cluster content: Original vs downsampled connectivity
Fowlkes-Mallows index
1
0.23
0
50
Remaining Connectivity [%]
100
L
Comparing morphology clustering vs downsampled connectivity clustering
Pearson correlation
1
0
50
Remaining Connectivity [%]
100
M
Fraction of iterations a DAN clusters with a group when all DANs have equal connections
1.2
Fraction of sampling iterations
0.0
γ5-10 γ5-5 γ5-6 γ5-1 γ5-2 γ5-4 γ5-12 γ5-13 γ5-7 γ5-16 γ5-21 γ5-19 γ5-20 γ5-23 γ5-25 γ5-24 β´2a-1 γ5-17 γ5-15 β´2a-2 β´2a-3 β´2a-4
N
Silhouette Method for input neuron clustering by morphology
Average silhouette width
0.40
0.24
*
2
Number of clusters k
38

**Figure S2 related to Figure 2**

(A-D) Related to Figures 2C and 2G. Representation of input neurons unique to (A) PAM-γ5 DANs (green), (B) PPL1-γ1pedc (red) and (C) PAM-β´2a DANs (blue). (D) Shared inputs (black). Whole brain outlined in grey. (E) Related to Figure 2D. Absolute (non-normalised) edge weight distribution between input neurons and DANs. The frequency at which an input to a given DAN with a certain number of connections occurs is shown as a function of the number of connections. DANs receive many inputs connected with only one or few synapses (skew to left). The long tailed distribution of PPL1-γ1pedc indicates it has some inputs with high absolute edge weights. (F) The Silhouette method was used to determine the appropriate cluster number (k) for connectivity clustering using Ward's criterion. Due to the high number of unique inputs with low connection number (Figure S2E), a single cluster for each DAN would be optimal. We chose k=7 to reflect the number of morphological DAN subclusters and the local maximum at 7 (asterisk) suggests k=7 is accurate. (G) Tanglegram illustrating comparison of connectivity clustering using Ward's criterion versus that from average linkage criterion. Identical clusters are formed although the between cluster relationship varies slightly. (H) Comparison of localization of postsynapses created by additional extensive review cycles compared to those identified following normal expert review. Marking the synapses along a γ5 DAN dendrite suggests both classes of synapses to be uniformly distributed. Plotting the distribution of synapses by measuring their positions as distance to the root of the dendritic tree confirms that both the standard and extensively reviewed populations are mixed along the dendrite. These analyses indicate that traced neurons that have undergone standard expert review, but not subjected to extensive review, can be considered to be randomly sampled. (I – L) To test whether input clustering is biased by an incompletely traced input network, we randomly down-sampled the connectivity dataset from 95% – 50% connectivity and compared the clustering obtained from these simulated datasets to the observed connectivity dataset (defined as 100% connectivity). (I) Plot illustrating correlation of connectivity distance matrices for differently down-sampled datasets versus our starting 100% connectivity dataset. The Pearson's correlation remains >0.9 even at 50% downsampling. The Null model based on randomised cluster allocation has a correlation of $4 \times 10^{-4}$. (J) To determine whether the DANs cluster in the same groups in the reduced (90%) connectivity dataset we compared their distributions in 10,000 iterations to those in the observed clusters at 100% connectivity. The fraction of times each neuron clusters with a particular group is shown, color coded according to the observed connectivity groups in Figure 2H (Also see cluster similarity analyses in Materials and Methods). Most DANs are reproducibly cluster with the same groups at 90% connectivity and only the γ5-19, γ5-20 and β´2a-2 show any propensity (<40%) to cluster with a different group. (K) The differences in clustering for 10,000 down-sampled datasets at each connectivity density can be individually compared to the observed 100% connectivity dataset using the Fowlkes-Mallows index (FMI). Even 50% connectivity datasets have an FMI (average = 0.71) well above the comparison for a randomised cluster allocation – the null model, FMI = 0.21. (L) Pearson's Correlation between connectivity and morphology for 100% and down-sampled connectivity datasets shows only subtle changes even at 50% connectivity, indicating that changes in (k) do not greatly alter the correlation between morphological and connectivity groups. (Randomised cluster allocation null model: -$6x10^{-4}$ ). (M) To test whether input connectivity clustering results were biased by the varying completeness of DAN post synapse annotation cluster similarity analyses were performed on 10,000 datasets in which all extensively reviewed PAM DANs had their connectivity (on average 216 inputs) down-sampled to the average connectivity of normally reviewed PAM DANs (on average 123 inputs). Only The γ5-20 and β´2a-2 DANs vary in their grouping with β´2a-2 likely to cluster with the γ5(uc) DANs and γ5-20 clustering alone. Clustering of all other DANs is stable indicating that the conclusions of this study do not result from biased completeness of tracing. (N) The Silhouette method was used to determine the accuracy of (k) for clustering based on input structure. Beyond 10 clusters the average silhouette width is high so taken with evaluation of cell body locations k = 20 was deemed to be accurate.

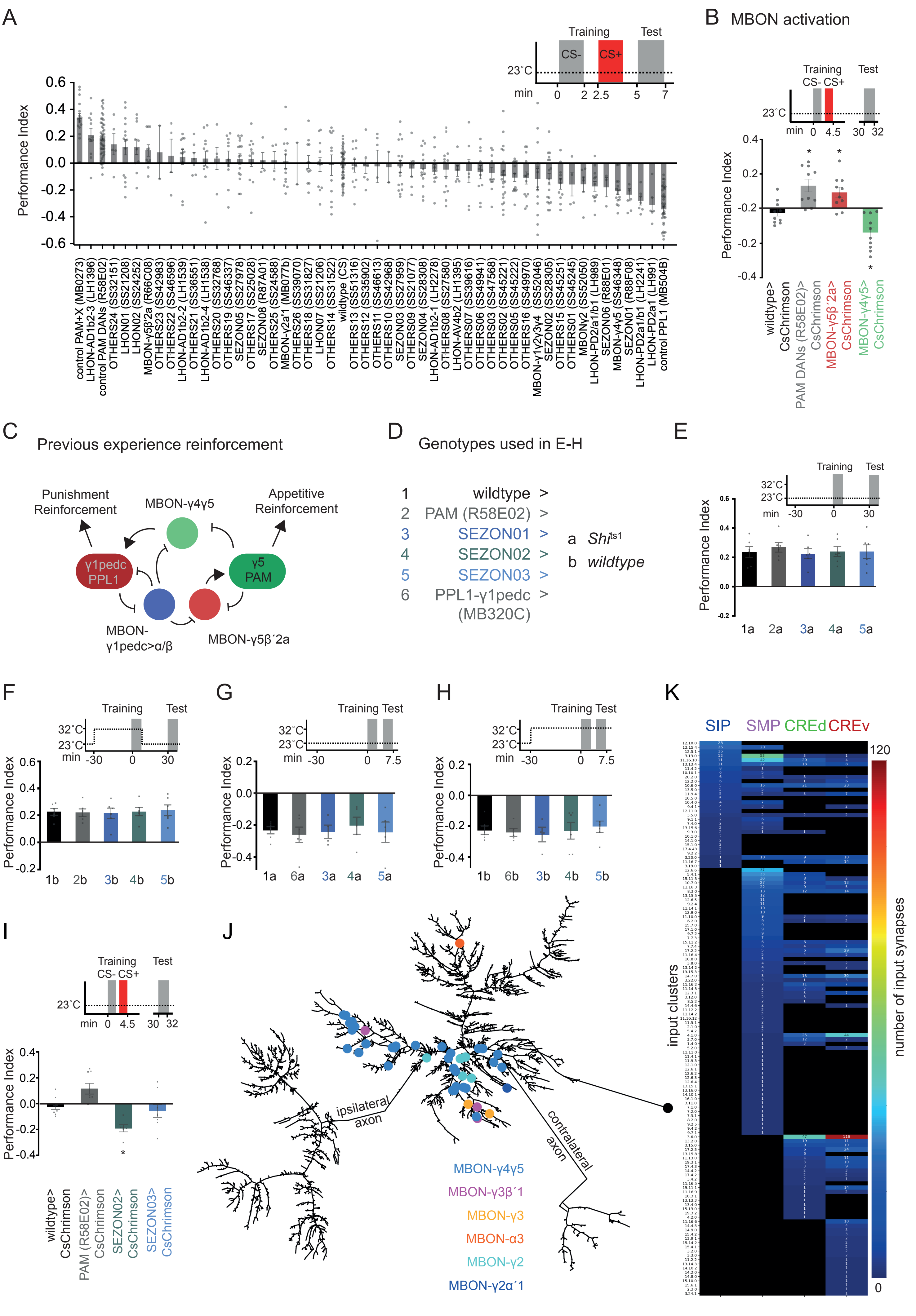

A
Training
Test
CS-
CS+
23˚C
min
0
2
2.5
5
7
Performance Index
0.6
0.4
0.2
0.0
-0.2
-0.4
-0.6
control PAM+X (MB0273)
LHON-AD1b2-3 (LH1396)
control PAM DANs (R58E02)
OTHERS24 (SS32151)
LHON01 (SS21208)
LHON02 (SS24252)
MBON-γ5β'2a (R66C08)
OTHERS23 (SS42983)
OTHERS22 (SS46596)
LHON-AD1b2-2 (LH1539)
OTHERS21 (SS36551)
LHON-AD1b2-4 (LH1538)
OTHERS20 (SS32768)
OTHERS19 (SS46337)
SEZON05 (SS27978)
OTHERS17 (SS25028)
SEZON08 (R87A01)
OTHERS25 (SS24588)
MBON-γ2a'1 (MB077b)
OTHERS26 (SS39070)
OTHERS18 (SS31827)
LHON07 (SS21206)
OTHERS14 (SS31522)
wildtype (CS)
OTHERS13 (SS51316)
OTHERS12 (SS35902)
OTHERS11 (SS46613)
OTHERS10 (SS42968)
SEZON03 (SS27959)
OTHERS09 (SS21077)
SEZON04 (SS28308)
LHON-AD1b2-1 (LH2278)
OTHERS08 (SS27580)
LHON-AV4b2 (LH1395)
OTHERS07 (SS39616)
OTHERS06 (SS49941)
OTHERS03 (SS47568)
OTHERS02 (SS45221)
OTHERS05 (SS45222)
OTHERS16 (SS49970)
MBON-γ1γ2γ3γ4 (SS52046)
SEZON02 (SS28305)
OTHERS15 (SS45251)
OTHERS01 (SS45245)
MBONγ2 (SS52050)
LHON-PD2/a1/b1 (LH989)
SEZON06 (R88E01)
MBON-γ4γ5 (SS46348)
SEZON01 (R88F08)
LHON-PD2a1/b1 (LH2241)
LHON-PD2a1 (LH991)
control PPL1 (MB504B)
B MBON activation
Training
CS- CS+
Test
23˚C
min 0 4.5 30 32
Performance Index
0.4
0.2
-0.2
-0.2
-0.4
wildtype> CsChrimson
PAM DANs (R58E02)> CsChrimson
MBON-γ5β´2a> CsChrimson
MBON-γ4γ5> CsChrimson
C
Previous experience reinforcement
Punishment Reinforcement
MBON-γ4γ5
Appetitive Reinforcement
γ1pedc PPL1
γ5 PAM
MBON-γ1pedc>α/β
MBON-γ5β´2a
D
Genotypes used in E-H
1 wildtype >
2 PAM (R58E02) >
3 SEZON01 >
4 SEZON02 >
5 SEZON03 >
6 PPL1-γ1pedc (MB320C) >
a Shi[ts1]
b wildtype
E
Training
Test
32˚C
23˚C
min -30 0 30
Performance Index
0.6
0.4
0.2
0.0
-0.2
1a 2a 3a 4a 5a
F
Training
Test
32˚C
23˚C
min -30 0 30
Performance Index
0.6
0.4
0.2
0.0
-0.2
1b 2b 3b 4b 5b
G
Training Test
32˚C
23˚C
min -30 0 7.5
Performance Index
0.2
0.0
-0.2
-0.4
1a 6a 3a 4a 5a
H
Training Test
32˚C
23˚C
min -30 0 7.5
Performance Index
0.2
0.0
-0.2
-0.4
1b 6b 3b 4b 5b
I
Training
CS- CS+
Test
23˚C
min 0 4.5 30 32
Performance Index
0.4
0.2
0.0
-0.2
-0.4
wildtype> CsChrimson
PAM (R58E02)> CsChrimson
SEZON02> CsChrimson
SEZON03> CsChrimson
J
ipsilateral axon
contralateral axon
MBON-γ4γ5
MBON-γ3β´1
MBON-γ3
MBON-α3
MBON-γ2
MBON-γ2α´1
K
SIP SMP CREd CREv
120
0
input clusters
number of input synapses

**Figure S3 related to Figure 3**

(A) Detail of screening results in Figure 3A. (B) MBON activation paired with odor exposure can write memories. Following identification in the screen, MBON-γ5β´2a and MBON-γ4γ5 were retested with 30 min memory being assessed. Activation of MBON-γ5β´2a formed weak positive memory ($p<0.0247$), whereas MBON-γ4γ5 activation formed aversive memory ($p<0.026$). PAM DANs ($p<0.0021$) (PAM DANs). Asterisks denote significance, one-way ANOVA, with Dunnett's post hoc test for multiple comparisons, n=10. (C) Previous experience reinforcement network for MBON-γ5β´2a and MBON-γ4γ5. Both MBONs receive dendritic input in the γ5 compartment but reinforce memories of opposite valence. MBON-γ4γ5 provides cross-compartmental feedback innervating PPL1-γ1pedc but not PAM-γ5 DANs. MBON-γ5β´2a provides feedback to the same compartment γ5 DANs and does not connect to PPL1-γ1pedc. MBON-γ1pedc>αβ (MVP2) is also part of the network and provides feed-forward inhibition [38]. (D) Key for groups shown in panels E-I. (E-I) Related to Figure 3 E and F. (E-H) Temperature and genetic controls for *Shi*$^{ts}$ block during training with sucrose (E and F) and DEET reinforcement (G and H). (I) SEZON activation paired with odor followed by testing 30 min memory replicates the screening phenotypes. SEZON02 produces aversive memories ($p<0.0245$, one-way ANOVA, with Dunnett's post hoc test for multiple comparisons, n=10). (J) Dendrogram of the dendrite of the PPL1-γ1pedc DAN showing upstream MBON presynapses (neato layout - graphviz). All MBON input goes to the CRE branches closest to the axon. MBON-γ4γ5 provides the strongest MBON input. (K) Relates to Figure 3G. Clustered input neurons provide selective input to specific branches of the PPL1-γ1pedc DAN dendritic tree.

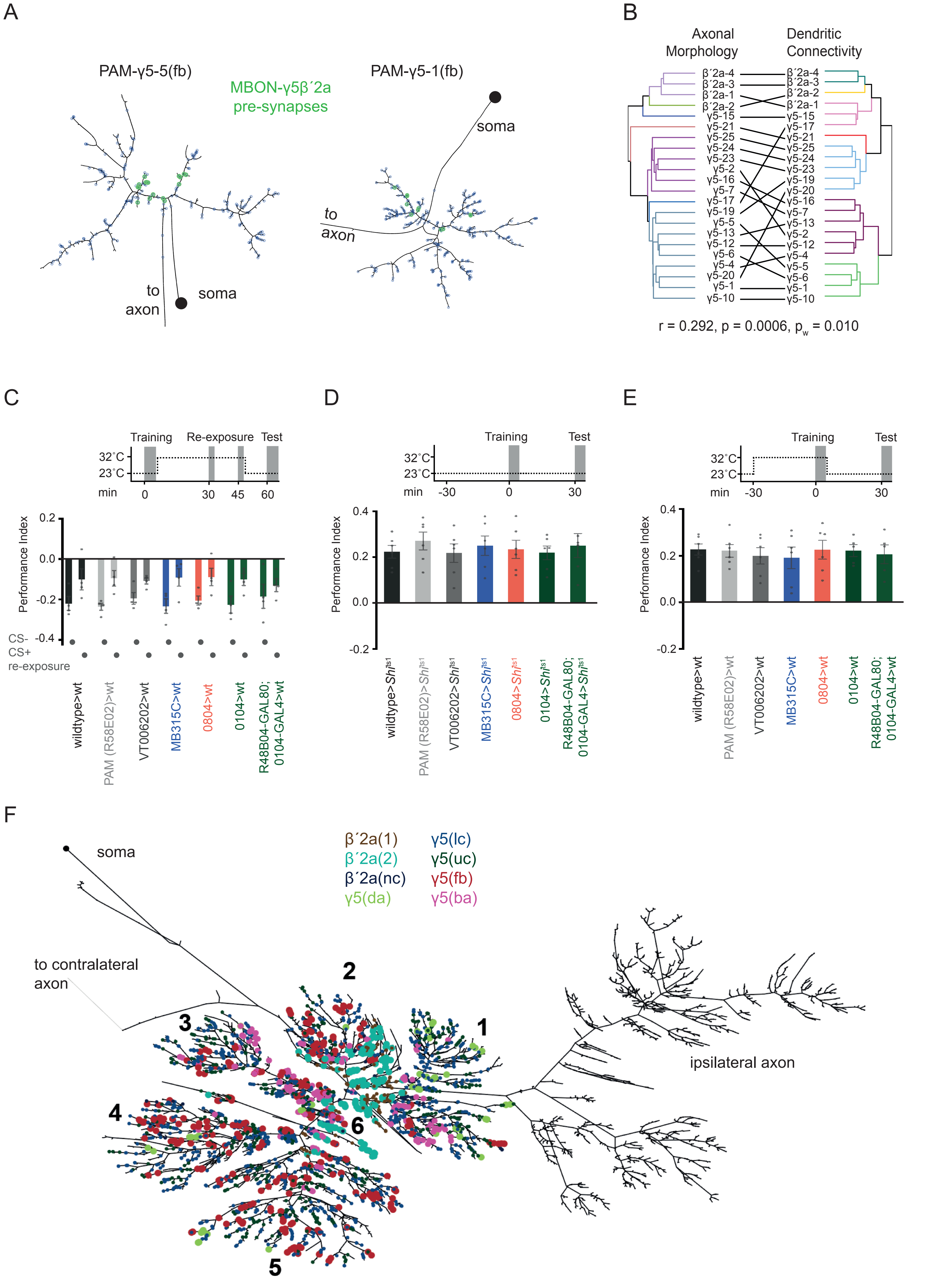
A
PAM-γ5-5(fb)
MBON-γ5β´2a pre-synapses
PAM-γ5-1(fb)
soma
to axon
B
Axonal Morphology
Dendritic Connectivity
r = 0.292, p = 0.0006, $p_w$ = 0.010
C
Training
Re-exposure
Test
32˚C
23˚C
min
Performance Index
CS-
CS+
re-exposure
wildtype>wt
PAM (R58E02)>wt
VT006202>wt
MB315C>wt
0804>wt
0104>wt
R48B04-GAL80; 0104-GAL4>wt
D
wildtype>$Shi^{ts1}$
PAM (R58E02)>$Shi^{ts1}$
VT006202>$Shi^{ts1}$
MB315C>$Shi^{ts1}$
0804>$Shi^{ts1}$
0104>$Shi^{ts1}$
R48B04-GAL80; 0104-GAL4>$Shi^{ts1}$
E
F
soma
β´2a(1)
β´2a(2)
β´2a(nc)
γ5(da)
γ5(lc)
γ5(uc)
γ5(fb)
γ5(ba)
to contralateral axon
ipsilateral axon

**Figure S4 related to Figure 4**

(A) Dendrograms of two PAM-γ5(fb) DANs with MBON-γ5β´2a synaptic inputs shown in green (neato layout - graphviz). MBON-γ5β´2a inputs localize close to the root of the dendrite (left) and on a single branch (right). MBON-γ5β´2a provides the largest fraction of the postsynaptic budget of γ5(fb) DANs contributed by a single neuron. (B) Tanglegram showing the correlation between morphological clustering of PAM DAN axons and dendritic connectivity clustering (Pearson's Correlation: $r=0.292$; $p=0.0006$, $p_w=0.01$ Mantel test). (C) Corresponds to Figure 4F. Heterozygous genetic driver controls show extinction learning (n=4). (D) Corresponds to Figure 4G. Permissive temperature control; no defects in 30 min sugar memory were observed (n=6). (E) Corresponds to Figure 4G. Genetic driver controls for 30 min sugar memory (n=5-6). (F) Dendrogram of MBON-γ5β´2a (neato layout - graphviz) with marked MBON postsynapses. Colors correspond to DAN morphology clusters providing the closest presynapse to the MBON postsynapses. We allowed a 2 µm radius around the respective postsynapse to account for dopamine diffusion and assign DAN presynapses. Dendritic branches are numbered 1-6 where 1, 2, and 6 are closest to the root of the dendrite. PAM-γ5(fb), -γ5(ba), -γ5(da), and -β´2a (2) DANs may influence specific branches, with the γ5(da) and γ5(fb) DANs connect exclusive branches and the γ5ba and β´2a (2) inputs localizing very close to the root of the dendrite. Morphologically distinct PAM DANs that receive different inputs might therefore modulate specific branches of the MBON-γ5β´2a dendrite.
