## Supplemental Data for "Input connectivity reveals additional heterogeneity of dopaminergic reinforcement in *Drosophila*"

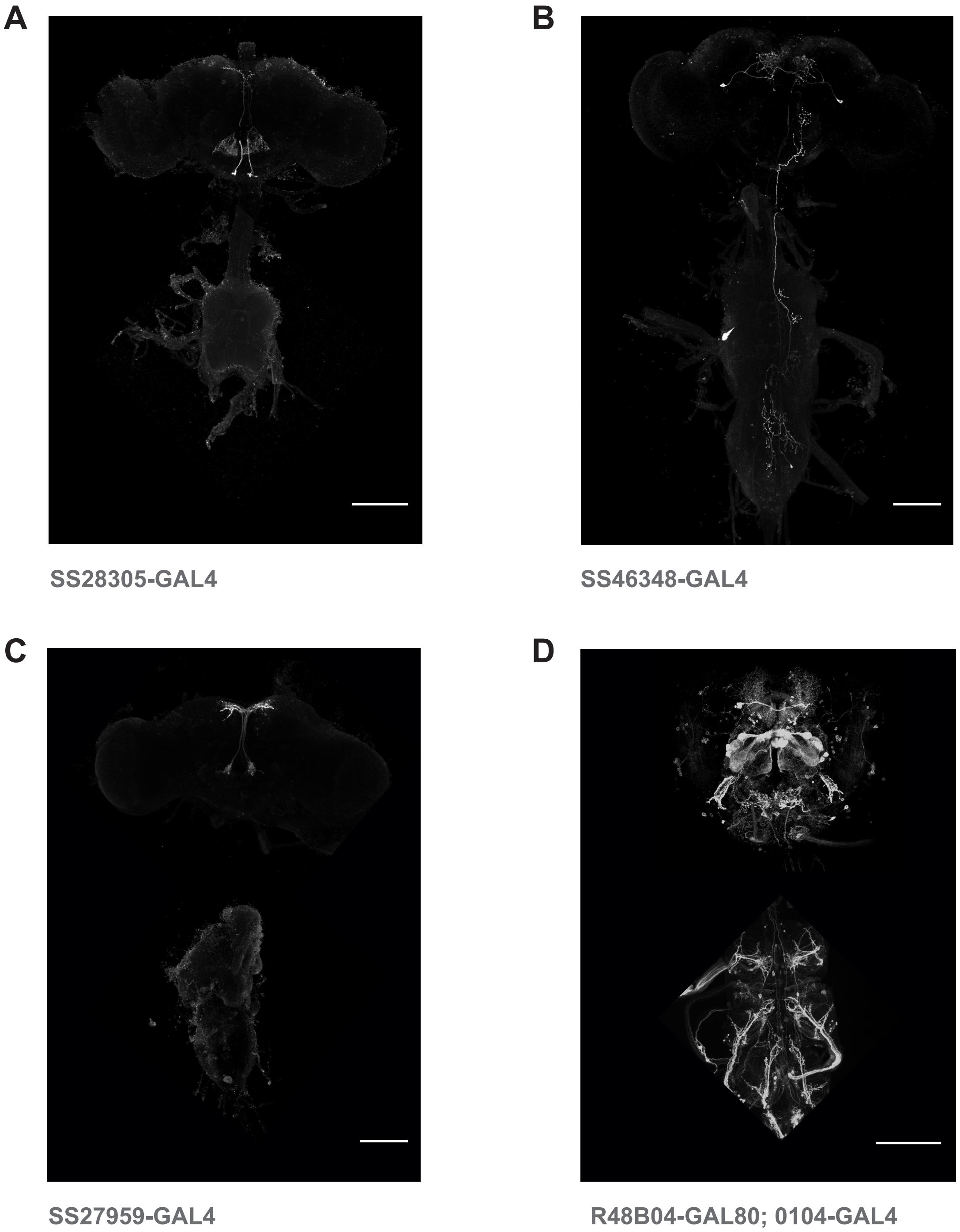


*Maximum intensity projections of confocal imaging stacks of Drosophila central and ventral nervous systems (CNS and VNC)*. Key GAL4 driver flies used in the functional experiments are shown expressing GFP (grey) in neurons of interest. Scalebar, 100µm. Due to COVID-19 enforced restrictions, we direct readers to the below listed publicly accessible databases for details/images of the other lines used in this study.

| Name | Collection | CNS and VNC |
|---|---|---|
| GMR58E02-GAL4 | FlyLight | For FlyLight and Vienna Tile lines: |
| GMR88F08 | FlyLight | https://www.janelia.org/project-team/flylight/tools-and-reagents |
| GMR66C08 | FlyLight | |
| MB320C-GAL4 | FlyLight | |
| MB315C-GAL4 | FlyLight | |
| VT006202-GAL4 | Vienna Tiles | For the InSite Collection: |
| 0104-GAL4 | InSite | http://www.columbia.edu/cu/insitedatabase/ |
| 0804-GAL4 | InSite | (W. Grueber and T. Clandinin Labs) |
